# Cryo-EM structure of the human Mixed Lineage Leukemia-1 complex bound to the nucleosome

**DOI:** 10.1101/737478

**Authors:** Sang Ho Park, Alex Ayoub, Young Tae Lee, Jing Xu, Hanseong Kim, Wei Zhang, Biao Zhang, Sojin An, Yang Zhang, Michael A. Cianfrocco, Min Su, Yali Dou, Uhn-Soo Cho

## Abstract

Mixed Lineage Leukemia (MLL) family histone methyltransferases are the key enzymes that deposit histone H3 Lys4 (K4) mono-/di-/tri-methylation and regulate gene expression in mammals. Despite extensive structural and biochemical studies, the molecular mechanism by which the MLL complexes recognize histone H3K4 within the nucleosome core particle (NCP) remains unclear. Here, we report the single-particle cryo-electron microscopy (cryo-EM) structure of the human MLL1 core complex bound to the NCP. The MLL1 core complex anchors on the NCP through RbBP5 and ASH2L, which interacts extensively with nucleosomal DNA as well as the surface close to histone H4 N-terminal tail. Concurrent interactions of RbBP5 and ASH2L with the NCP uniquely align the catalytic MLL1^SET^ domain at the nucleosome dyad, allowing symmetrical access to both H3K4 substrates within the NCP. Our study sheds light on how the MLL1 complex engages chromatin and how chromatin binding promotes MLL1 tri-methylation activity.

## INTRODUCTION

The nucleosome core particle (NCP), consisting of an octameric core of histone proteins (two of each H2A, H2B, H3, and H4) and 146 base-pairs of genomic DNA, represents the first level of eukaryotic DNA packaging (Kornberg and Thomas, 1974). It is further organized into higher order chromatin structures. Cell specific transcription program, in large part, is governed by chromatin accessibility, which is actively regulated by histone modifying enzymes and ATP-dependent chromatin remodeling complexes. In recent years, X-ray crystallography and single-particle cryo-EM studies have shed light on how these chromatin-associating complexes interact with the NCP for respective physiological functions. Most, if not all, chromatin complexes engage the ‘acidic-patch’ region of the NCP through variations of an ‘arginine-finger’ motif (Armache et al., 2011; Barbera et al., 2006; Kato et al., 2013; Makde et al., 2010), highlighting common features among chromatin interacting protein complexes. It remains unclear whether the recognition mode of the NCP is universal for chromatin interacting complexes.

Among histone post-translational modifications, the states of histone H3 lysine4 methylation (H3K4me) (i.e., mono-, di-, tri-methylation) are exquisitely modulated at important DNA regulatory regions including active gene promoters, gene bodies and distal regulatory enhancers (Rao and Dou, 2015). In particular, H3K4me3 is highly correlative to the transcriptionally active- and open-chromatin regions (Chen et al., 2015; Ruthenburg et al., 2007) and is shown to actively recruit the basic transcription machinery, ATP-dependent chromatin remodeling complexes, and histone acetyltransferases (Lauberth et al., 2013; Ruthenburg et al., 2007; Taverna et al., 2007; Vermeulen et al., 2007; Wysocka et al., 2006). In contrast, H3K4me1 is a prevalent mark often found at poised or active distal enhancers (Herz et al., 2012). Specific regulation of the H3K4me states may play a critical role in important physiological processes in cells.

The mixed lineage leukemia (MLL) family enzymes, including MLL1-4/KMT2A-2D, SET1A/KMT2F and SET1B/KMT2G), are the major histone lysine 4 (K4) methyltransferases in mammals. They contain an evolutionarily-conserved catalytic <u>S</u>u(Var)3-9, <u>E</u>nhancer of Zeste, <u>T</u>rithorax (SET) domain (Rea et al., 2000). Biochemical studies, by us and others, have shown that the SET domain stably interacts with four highly conserved proteins, i.e. RbBP5 (<u>r</u>etino<u>b</u>lastoma-<u>b</u>inding <u>p</u>rotein 5), ASH2L (<u>A</u>bsent, <u>s</u>mall, <u>h</u>omeotic disks-2-like), WDR5 (<u>WD</u>40 <u>r</u>epeat-containing protein 5) and DPY30 (<u>D</u>um<u>PY</u> protein 30) (Dou et al., 2006)(van Nuland et al., 2013). The MLL1 core components are able to increases the MLL1^SET^ activity on mono- and di-methylation of histone H3K4 by ∼600 fold (Patel et al., 2009). The molecular mechanism by which MLL1 core components stimulate MLL1^SET^ activity has been elegantly demonstrated by a series of structural studies including the human MLL1/3^SET^-ASH2L^SPRY^-RbBP5^330-375^ subcomplex (Li et al., 2016), the homologous yeast SET1 complexes (Hsu et al., 2018; Qu et al., 2018) as well as individual mammalian core components (Avdic et al., 2011; Cosgrove and Patel, 2010; Rao and Dou, 2015). However, it is still unclear how tri-methylation activity of MLL1, which has been widely-reported *in vivo* (Dou et al., 2005; Guenther et al., 2005; Katada and Sassone-Corsi, 2010; Wysocka et al., 2005), is regulated. Up to date, the structures of the MLL family enzymes are determined with either no H3 or H3 peptide as the substrate. It remains unclear how the MLL1 complex binds and catalyzes H3K4 methylation on the NCP and more importantly, how MLL1 activity, especially the tri-methylation activity, is regulated on chromatin.

Mutations of MLL proteins have been widely reported in a variety of congenital human syndromes including Kabuki (Hannibal et al., 2011; Kluijt et al., 2000; Li et al., 2011; Micale et al., 2011; Ng et al., 2010; Paulussen et al., 2011), Wiedemann-Steiner (Jones et al., 2012; Mendelsohn et al., 2014; Strom et al., 2014), and Kleefstra spectrum syndromes (Strom et al., 2014) as well as a wide spectrum of human malignancies (Rao and Dou, 2015). Similarly, aberrant expression and recurrent mis-sense mutations of ASH2L and RbBP5 have also been identified in human cancers of different origins (Bochynska et al., 2018; Butler et al., 2017; Ge et al., 2016; Magerl et al., 2010). It is important to understand how MLL activity on chromatin is modulated during disease development.

Here, we report the single-particle cryo-EM structure of the human MLL1 core complex bound to the NCP. It not only reveals the overall architecture of the human MLL1 complex with full-length core components, but also illustrates how the MLL1 core complex engages the chromatin. Importantly, we show that the MLL1 core complex docks on the NCP through concurrent interactions of ASH2L/RbBP5 with nucleosomal DNA and histone H4. This unique configuration aligns the catalytic MLL1^SET^ domain at the nucleosome dyad, which allows the symmetric access to both H3K4 substrates. Our structure sheds new light on how the MLL1 complex binds to the chromatin and how its activity for H3K4me3 is regulated.

## RESULTS

### Architecture of the Human MLL1 Core Complex Bound to the NCP

A recombinant human MLL1 core complex (MLL1^RWSAD^) containing <u>R</u>bBP5 (residues 1–538), <u>W</u>DR5 (residues 25–330), MLL1^<u>S</u>ET^ (residues 3762–3969), <u>A</u>SH2L (residues 1– 534) and <u>D</u>PY30 (residues 1–99) was reconstituted *in vitro* (Figures 1A and S1A). An electrophoretic mobility shift assay demonstrated that MLL1^RWSAD^ binds to the NCP with modest affinity (Figures S1B-C). We found that this complex showed markedly enhanced activity for higher methylation states (i.e., H3K4me2 and H3K4me3) when the NCP was used as a substrate (Figure 1B). To understand the underlying mechanism, we determined the single-particle cryo-EM structure of reconstituted MLL1^RWSAD^ bound to the recombinant NCP. The cryo-EM structure of MLL1^RWSAD^-NCP was determined at a 6.2 Å resolution (Figures 1C, S2, and S3A and Table 1). The composite map of MLL1^RWSAD^-NCP was generated after local filtering to the estimated resolution to avoid over-interpretation (Figures 1C and S2). In parallel, the cryo-EM maps of RbBP5-NCP and <u>R</u>bBP5-<u>W</u>DR5-MLL1^<u>S</u>ET^ (MLL1^RWS^)-NCP subcomplexes were derived from the MLL1^RWSAD^-NCP dataset, and reconstructed at 4.2 Å and 4.5 Å resolution, respectively (see the STAR methods; Figures S2 and S3B-D). The model structure of the MLL1^RWSAD^-NCP complex (Figure 1D) was built by rigid-body fitting and real space refinement using previously published crystal structures of mouse RbBP5 (PDB ID: 5OV3) (Mittal et al., 2018), human WDR5 (PDB ID: 2H14) (Couture et al., 2006), human MLL1^SET^-ASH2L^SPRY^-RbBP5^330-375^ (PDB ID: 5F6L) (Li et al., 2016), a DPY30 dimer (PDB ID: 6E2H) (Haddad et al., 2018), and the 601-NCP (PDB ID: 3MVD) (Makde et al., 2010).

**Table 1.** Cryo-EM Data Collection, Refinement, and Validation Statistics.

|  | MLL1<sup>RWSAD</sup>-NCP<br>(EMD-20512)<br>(PDB: 6PWV) | MLL1<sup>RWS</sup>-NCP<br>(EMD-20513)<br>(PDB: 6PWW) | RbBP5-NCP<br>(EMD-20514)<br>(PDB: 6PWX) |
| --- | --- | --- | --- |
| **Data Collection and Processing** |  |  |  |
| Magnification | 29,000 |  |  |
| Voltage (kV) | 300 |  |  |
| Electron exposure (e-/Å<sup>2</sup>) | 64 |  |  |
| Defocus range (μm) | -1.5 to -3.5 |  |  |
| Pixel size (Å) | 1.01 |  |  |
| Symmetry imposed | C1 |  |  |
| Initial particle images (no.) | 712,198 |  |  |
| Final particle images (no.) | 8,433 | 21,114 | 32,563 |
| Map resolution (Å) | 6.2 | 4.5 | 4.2 |
| FSC threshold | 0.143 | 0.143 | 0.143 |
| **Refinement** |  |  |  |
| Initial model used (PDB code) | 3MVD, 5OV3,<br>2H14, 5F6L, 6E2H | 3MVD, 5OV3,<br>2H14, 5F6L | 3MVD, 5OV3 |
| Model resolution (Å) | 6.7 | 4.7 | 4.3 |
| FSC threshold | 0.5 | 0.5 | 0.5 |
| Map sharpening *B* factor (Å<sup>2</sup>) | -189 | -157 | -100 |
| **Model composition** |  |  |  |
| Non-hydrogen atoms | 21,781 | 18,174 | 14,411 |
| Protein residues | 2,005 | 1,550 | 1,074 |
| Nucleotides | 292 | 292 | 292 |
| Ligands | 2 | 2 | - |
| ***B* factor (Å<sup>2</sup>)** |  |  |  |
| Protein | 192.99 | 109.71 | 107.65 |
| Nucleotide | 48.20 | 36.73 | 20.20 |
| Ligand | 781.00 | 826.18 | - |
| **Rmsds** |  |  |  |
| Bond lengths (Å) | 0.005 | 0.004 | 0.003 |
| Bond angles (°) | 0.688 | 0.612 | 0.579 |
| **Validation** |  |  |  |
| MolProbity score | 2.54 | 2.21 | 2.38 |
| Clashscore | 31.16 | 21.65 | 13.50 |
| Poor rotamers (%) | 1.42 | 1.22 | 3.58 |
| **Ramachandran plot** |  |  |  |
| Favored (%) | 93.05 | 95.54 | 95.45 |
| Allowed (%) | 6.95 | 4.46 | 4.55 |
| Disallowed (%) | 0 | 0 | 0 |

**Figure 1.**
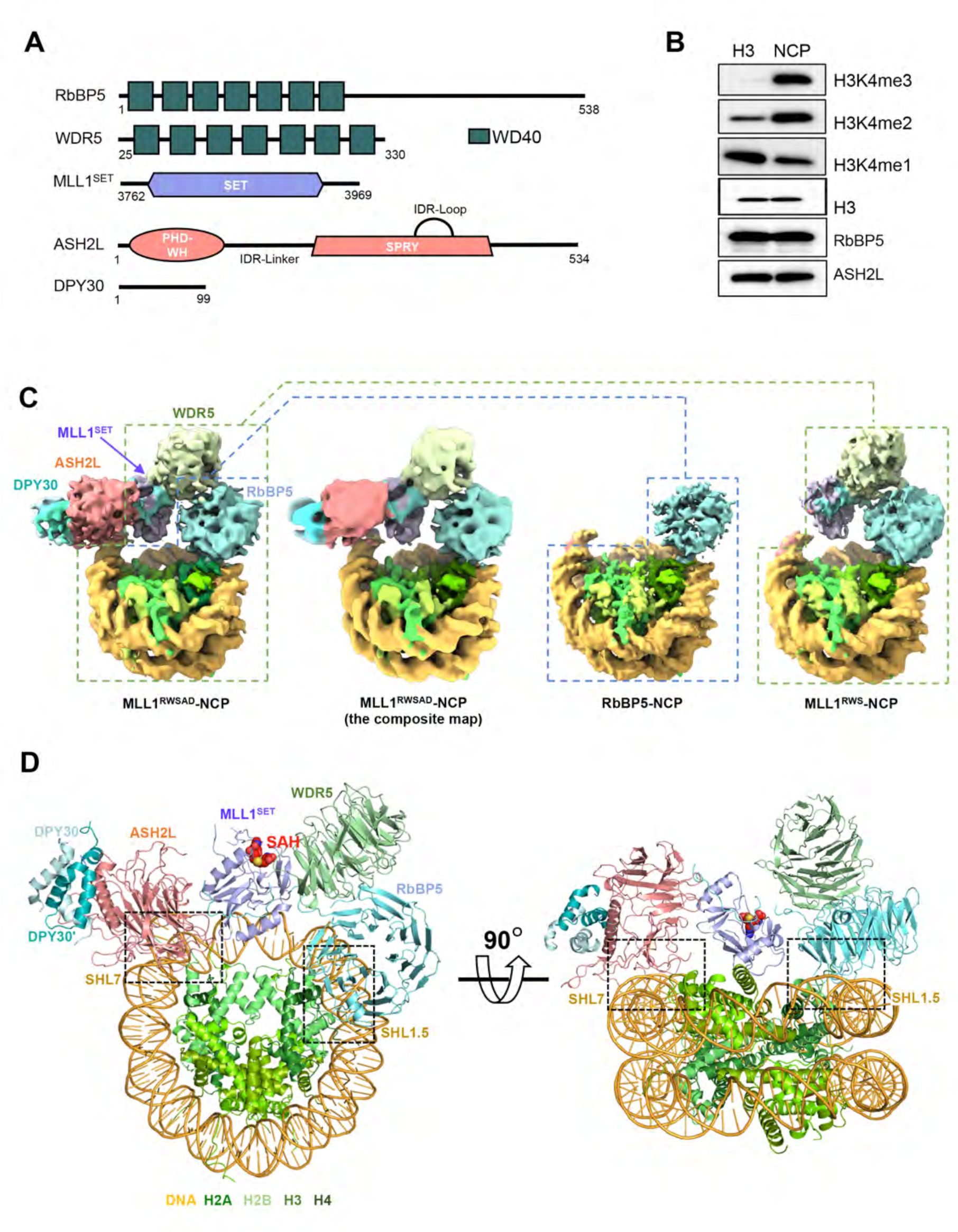
Cryo-EM Structure of the MLL1^RWSAD^-NCP Complex. (A) Schematic domain architectures for the core components of the human MLL1 complex used in the cryo-EM study. (B) Immunoblot to detect H3K4 methylation in the *in vitro* histone methyltransferase assay. Antibodies indicated on right. The substrates were free recombinant histone H3 (left) and the NCP (right), respectively. Immunoblots for unmodified H3 as well as RbBP5 and ASH2L included as controls. (C) Cryo-EM 3D reconstruction of the MLL1^RWSAD^-NCP complex. The composite map of MLL1^RWSAD^-NCP was locally filtered to the estimated resolution. The subcomplexes, i.e., RbBP5-NCP and MLL1^RWS^-NCP, shown in dashed boxes. (D) Top (left) and front (right) views of the MLL1^RWSAD^-NCP structure. The S-adenosyl-L-homocysteine (SAH) was represented as a sphere (red) and the MLL1 core components shown in cartoon representation (RbBP5: cyan, WDR5: green, MLL1^SET^: slate, ASH2L: orange, and DPY30 dimer: cerulean and teal). The 147 base-pair Widom 601 DNA and four histones were colored as indicated on bottom. Two black dashed squares highlighted the nucleosome contact points near SHL1.5 and SHL7 by MLL1^RWSAD^. Illustrations of the protein structure and cryo-EM maps used in all figures were generated with PyMOL (Delano Scientific, LLC) and Chimera (Pettersen et al., 2004)/ ChimeraX (Goddard et al., 2018).

The overall architecture of the MLL1^RWSAD^-NCP complex showed that MLL1^RWSAD^ anchors at the edge of the NCP through two core components, RbBP5 and ASH2L simultaneously (Figure 1D). In the NCP, DNA <u>s</u>uper<u>h</u>elical location 7 (SHL7) and SHL1.5, together with H4 N-terminal tail, were involved in the interaction with MLL1^RWSAD^ (Figure 1D). Notably, domains in the MLL1^RWSAD^-NCP complex were dynamically associated with each other and showed multiple conformations (Figure S3E). However, the overall architecture was conserved in all sub-classes of the MLL1^RWSAD^-NCP structures (Figure S3E). Distinct from many of previously reported NCP-recognizing protein or protein complexes (Zhou et al., 2019), the MLL1 core complex did not interact with the acidic patch region of the NCP.

### RbBP5 Binds the NCP through Both DNA and Histone H4 Tail Recognition

In the MLL1^RWSAD^-NCP complex, the RbBP5-NCP interfaces were less dynamic. The sub-population particles of RbBP5-NCP from the MLL1^RWSAD^-NCP dataset were resolved at 4.2 Å resolution (Figures 2A, S2, and S3B). The regions of mouse RbBP5 in model fitting shared 100% sequence identity with human RbBP5 (Figure S4A). The structure showed that RbBP5 bound to the NCP by simultaneously engaging DNA (SHL1.5) and histone H4 N-terminal tail. The interactions involved six consecutive loops emanating from the WD40 repeats of RbBP5 (Figure 2A). Characteristic features of RbBP5 (e.g. unique helix, anchoring loop, and insertion loop) were well matched into the cryo-EM map of RbBP5-NCP subcomplex (Figure S4B). Notably, RbBP5 interacted with DNA SHL1.5 through four positively-charged arginine residues (Quad-R) located at β18–β19 (R220), β20–β21 (R251), β22–β23 (R272), and β24–β25 (R294) loops, respectively (Figure 2B). The Quad-R participated in electrostatic interactions with the DNA phosphate backbone (Figure 2B). Disruption of RbBP5-NCP interaction significantly reduced the activity of the MLL1 core complex. Mutations of the Quad-R residues to alanine (A) led to reduction of H3K4me3 and to a lesser degree, H3K4me2 (Figure 2C). The effect was more drastic when Quad-R residues were mutated to glutamic acid (E) (Figure 2D). Systematic alteration of three, two or one arginine residue(s) in Quad-R showed that at least two arginine residues were required for optimal H3K4me3 activity (Figure 2C).

**Figure 2.**
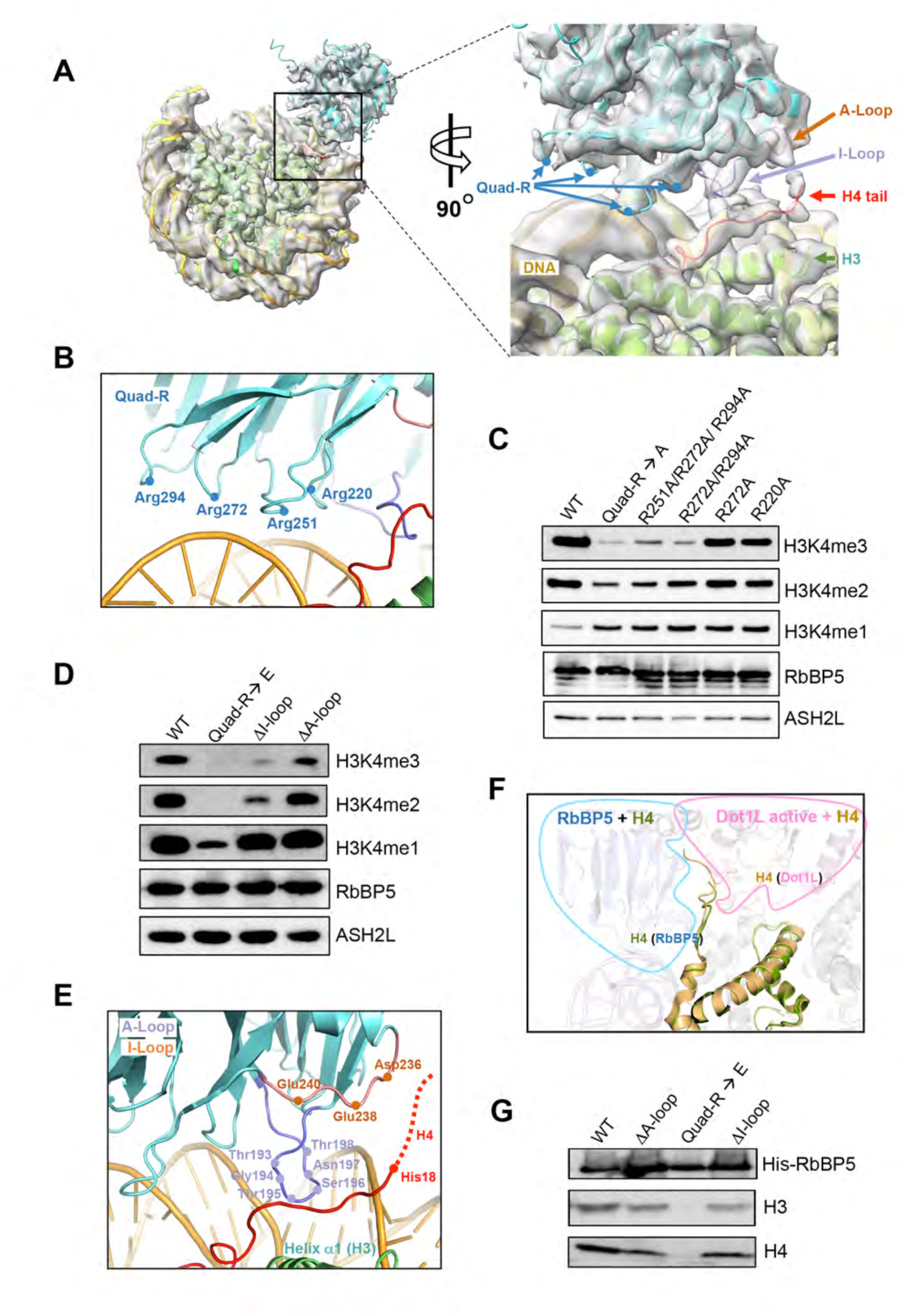
RbBP5 Interaction with the NCP. (A) The cryo-EM structure of the RbBP5-NCP subcomplex (4.2 Å). The interaction interface was enlarged and shown on right. Insertion (I)-loop, Anchoring (A)-loop, and Quad-R of RbBP5 as well as the H4 tail highlighted in orange, purple, blue and red, respectively. Histone H3 shown in green. (B) Interaction of Quad-R, as indicated, with DNA backbone. Red line, histone H4 tail. (C) Immunoblot to detect *in vitro* histone methyltransferase activity with the NCP as the substrate. The MLL1^RWSAD^ complex reconstituted with wild type and Quad-R mutated RbBP5 indicated on top. (D) Immunoblot to detect *in vitro* histone methyltransferase activity with the NCP as the substrate. The MLL1^RWSAD^ complex reconstituted with RbBP5 wild type and deletion mutant proteins indicated on top. (E) The interface between RbBP5 and the H4 tail. Key residues on RbBP5 I-/A-loops indicated. The H4 tail (His18 to core) represented by a red line and the extended tail beyond His18 represented by a dash line. (F) Structural superposition of the H4 tails upon RbBP5 (green) and Dot1L (PDB ID: 6NJ9) (Worden et al., 2019) binding. The RbBP5 and Dot1L at the interfaces enclosed by the blue and pink outlines, respectively. (G) *In vitro* pull-down assay for RbBP5 and the NCP. Ni-NTA-bound fractions were shown and His-tagged wild type or mutant RbBP5 proteins shown on top. Immunoblot for H3 used to detect the NCP in the bound fraction. Immunoblot for H4 used as a control.

The second RbBP5-NCP interface includes two loops, an insertion loop (β16–β17 loop, referred to herein as I-loop) and an anchoring loop (β19–β20 loop, referred to herein as A-loop), of RbBP5 (Figures 2A, 2E, S4B, and S5A). Both I- and A-loops are evolutionarily conserved in higher eukaryotes (Figure S5B). The I-loop was positioned between the N-terminal tail of histone H4 and nucleosomal DNA (Figure 2E). The A-loop run parallel to H4 tail, which was positioned between the I-/A-loops of RbBP5 and the helix α1 (Leu65–Asp77) of histone H3 (Figure 2E). This H4 tail-mediated nucleosome recognition of RbBP5 resembles that of the active-state DOT1L (Anderson et al., 2019; Worden et al., 2019) (Figure 2F). Similar to Quad-R, deletion of I-loop and to a lesser degree A-loop, reduced the activity of the MLL1 core complex for H3K4me3 and H3K4me2 (Figure 2D). Importantly, RbBP5-NCP interaction is specifically required for MLL1 activity on the NCP (Figure 2D). Mutations in Quad-R, I-loop, and A-loop had no effects on mono-, di-, tri-methylation of free H3 (Figure S5C). Among RbBP5-NCP interactions, Quad-R was the main contributor to NCP binding. The mutation of Quad-R significantly reduced RbBP5 binding to the NCP, while the deletion of I- and A-loops only modestly affected NCP binding (Figure 2G).

### Structural Organization of WDR5, MLL1^SET^, and ASH2L^SPRY^ Sub-complex

To resolve the structural organization of WDR5, MLL1^SET^, and ASH2L^SPRY^ sub-complex, we reconstructed the MLL1^RWS^-NCP subcomplex (Figure 1C) (Figure S2 and the STAR methods) and successfully docked the crystal structures of human WDR5 (PDB ID: 2H14) (Couture et al., 2006) and MLL1^SET^-ASH2L^SPRY^-RbBP5^330-375^ (PDB ID: 5F6L) (Li et al., 2016) into the cryo-EM maps (Figure 3A and 3B). The secondary structural components of MLL1^SET^ (Southall et al., 2009), including α–helices and β-hairpin of the SET-I, SET-N, and SET-C domains (dotted circles), fitted well into the cryo-EM map (Figure 3B). Similar to MLL1^SET^, distinctive features of WDR5 and ASH2L^SPRY^ were also well-defined in the cryo-EM structure (Figure 3A and 3B). Importantly, our structure indicated that the WDR5-MLL1^SET^-ASH2L^SPRY^ sub-complex did not make direct contacts with nucleosomal DNA, which was experimentally confirmed by the gel mobility assays (Figure S5D and data not shown). The catalytic site of the MLL1^SET^ domain was pointing outward, which might confer distance restraint on substrate accessibility (see below). Furthermore, the overall domain architecture of the MLL1 core complex within MLL1^RWSAD^-NCP was largely conserved in the NCP-free yeast SET1 complexes (Hsu et al., 2018; Qu et al., 2018), suggesting that NCP binding may not require or induce major conformational changes in the MLL1 core complex (Figure 3C).

**Figure 3.**
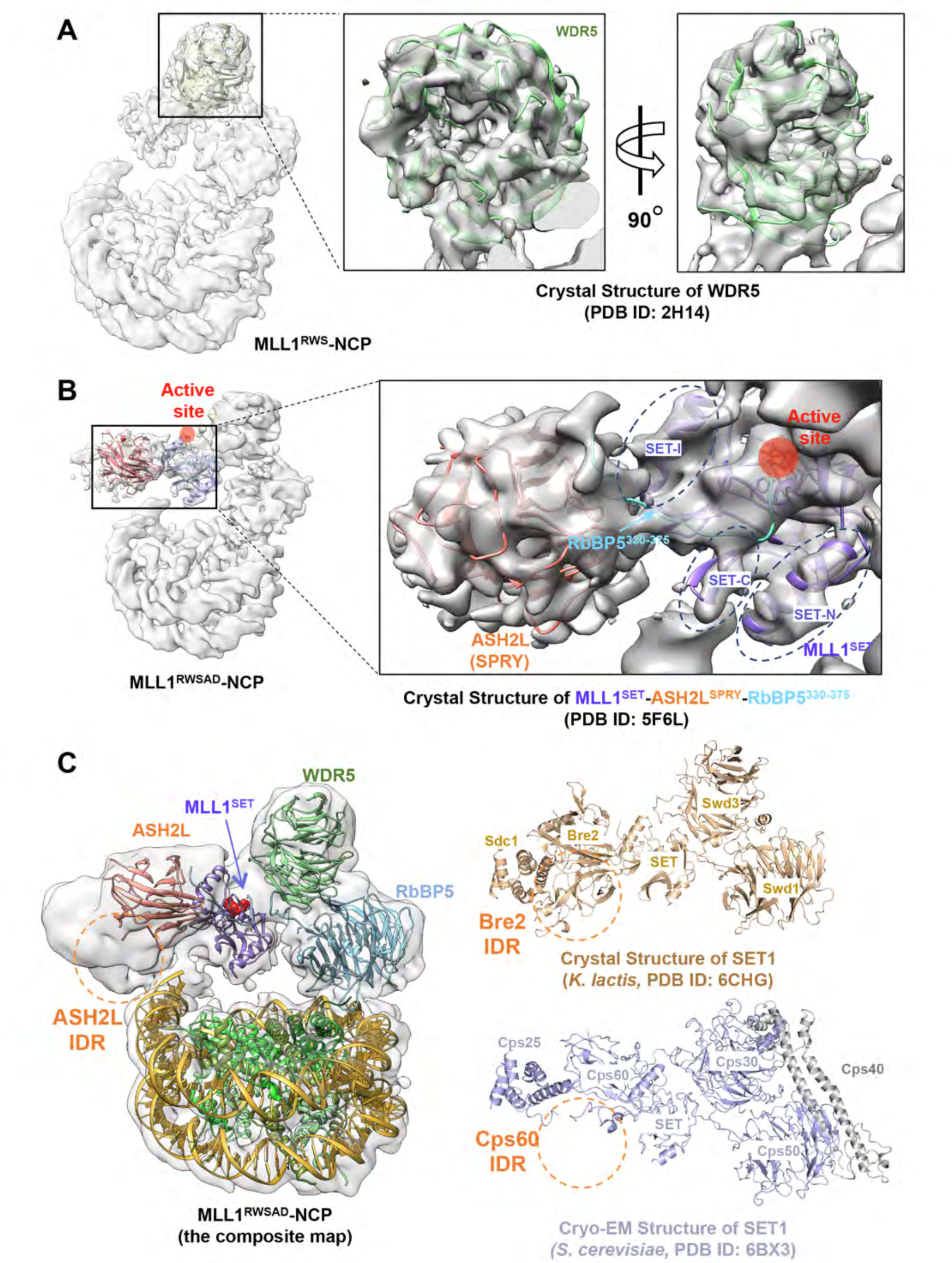
The WDR5-MLL1^SET^-ASH2L^SPRY^-NCP Subcomplex. (A) Rigid-body fitting of human WDR5 crystal structure (PDB ID: 2H14) (Couture et al., 2006) into the cryo-EM map of MLL1^RWS^-NCP. Secondary structures of WDR5 were shown in green. (B) Rigid-body fitting of MLL1^SET^ and ASH2L^SPRY^ into the Cryo-EM map of MLL1^RWSAD^-NCP. The MLL1^SET^-RbBP5^330-375^-ASH2L^SPRY^ crystal structure (PDB ID: 5F6L) (Li et al., 2016) was used. Characteristic secondary structures of MLL1^SET^ (SET-I, SET-C and SET-N) were shown within black dashed circles. The catalytic active site represented by red sphere. (C) Comparison of human MLL1-NCP structure with yeast SET1 crystal structure (*K. lactis*, dark goldenrod, PDB ID: 6CHG, right top) (Hsu et al., 2018) and cryo-EM structure (*S. cerevisiae*, light slate blue, PDB ID: 6BX3, right bottom) (Qu et al., 2018). Orange circles indicated IDR regions of ASH2L (human), Bre2 (*K. lactis*), and Cps60 (*S. cerevisiae*). An extra domain in *S. cerevisiae* SET1 complex, Cps40, colored grey.

### Dynamic ASH2L-NCP Interaction is Critical for H3K4me3

The second docking point of the MLL1 core complex to the NCP was provided by the intrinsically disordered regions (IDRs) of ASH2L (Figures 1D and 3C). The ASH2L-NCP interface was highly dynamic in solution (Figure S3E), making it challenging to visualize the molecular details. Similar dynamic behaviour was observed for IDR of the yeast homologue Cps60 (Figure 3C), which was not resolved in the cryo-EM structure of the yeast SET1 complex (Qu et al., 2018). Since the crystal structure of full-length human ASH2L has not been reported, we employed the protein structure prediction approach using the iterative template-based fragment <u>ass</u>embly <u>r</u>efinement (I-TASSER) method (Roy et al., 2010; Zhang, 2008). The crystal structure of yeast Bre2 was used as template (PDB ID: 6CHG) (Hsu et al., 2018) to build the ASH2L plant homeodomain-wing helix (PHD-WH)/IDRs model (Figures 4A and S6A). After resolving minor clashes, we were able to reliably dock ASH2L IDRs into the cryo-EM map of MLL1^RWSAD^-NCP (Figures 4A, S6A, and S6B). The model of MLL1^RWSAD^-NCP showed that ASH2L IDRs interacted with SHL7 of nucleosomal DNA (Figures 1D and 4B). Surprisingly, the PHD-WH domain of ASH2L was located outside of the cryo-EM map (Figure S7A) despite the reported function in DNA binding (Chen et al., 2011; Sarvan et al., 2011).

**Figure 4.**
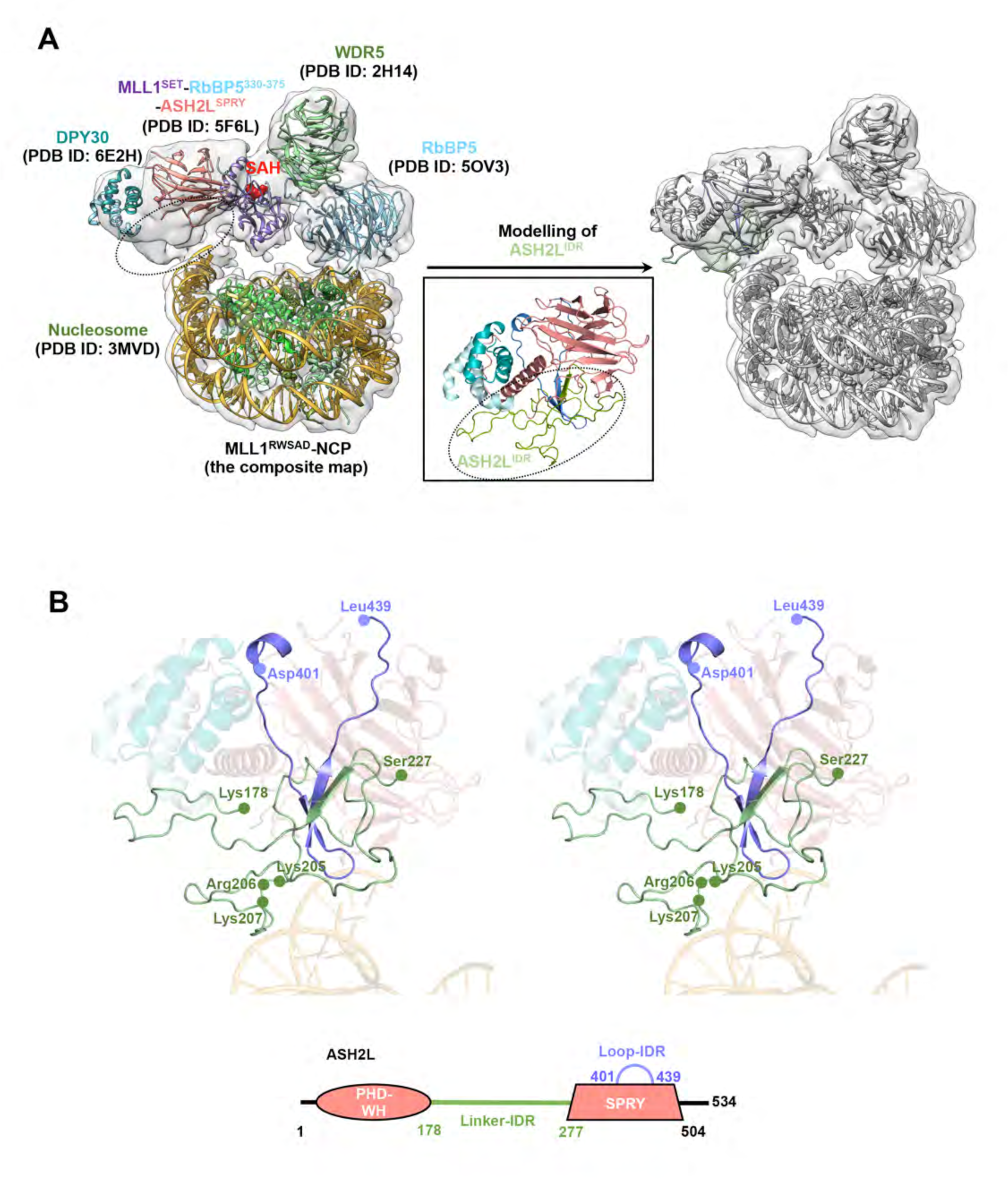
ASH2L Interacts with the Nucleosomal DNA through IDRs. (A) Structure prediction of ASH2L^IDR^. The structure of ASH2L IDR regions was not available and thus not assigned in the corresponding cryo-EM map (dashed circle). The structure prediction approach was employed to model ASH2L IDR regions as described in the STAR methods. Linker-IDR colored green and Loop-IDR colored blue in the ASH2L^IDR^ model structure. (B) Stereo-view of the ASH2L-DPY30 structure and DNA contacts. The structure of ASH2L using the crystal structure of ASH2L^SPRY^ (PDB ID: 5F6L) (Li et al., 2016) and the model structure of ASH2L^IDR^. Structures of ASH2L^SPRY^/DPY30/DNA shown as transparent and key residues in ASH2L IDR were labeled. The domain architecture of full-length ASH2L shown at the bottom.

Our MLL1^RWSAD^-NCP model pinpointed a short stretch of positively-charged residues (i.e., K205/R206/K207) in the ASH2L Linker-IDR to make contacts with nucleosomal DNA (Figure 4B). These positively-charged residues were highly conserved in ASH2L homologs in higher eukaryotes (Figure 5A). To biochemically validate the model structure of ASH2L IDRs, we first examined the importance of key residues involved in the NCP interaction. As shown in Figure 5B, ASH2L directly interacted with the NCP, resulting in a mobility shift in the native gel. However, deletion of both PHD-WH (residues 1–178) and Linker-IDR (residues 178–277), but not PHD-WH alone, abolished ASH2L interaction with the NCP (Figure 5B). Further truncation of ASH2L Linker-IDR identified that residues 202–207 were important for NCP interaction, consistent with our ASH2L model (Figure 4B). Binding of ASH2L to the NCP was critical for MLL1 methyltransferase activity. Deletion of ASH2L Linker-IDR completely abolished the MLL1 activity on the NCP (Figure 5C, left). Similarly, deletion of ASH2L residues 202–207 or mutations of residues K205/R206/K207 to alanine significantly reduced MLL1 activity on the NCP for H3K4me3 (Figure 5C, right), but not on free H3 (Figure S7B). This result, together with that of RbBP5, suggests that MLL1-NCP interactions specifically promote tri-methylation of H3K4. Notably, deletion of ASH2L Linker-IDR led to more drastic reduction of overall H3K4me, suggesting additional mechanisms by which Linker-IDRs may contribute to MLL1 regulation (see discussion).

**Figure 5.**
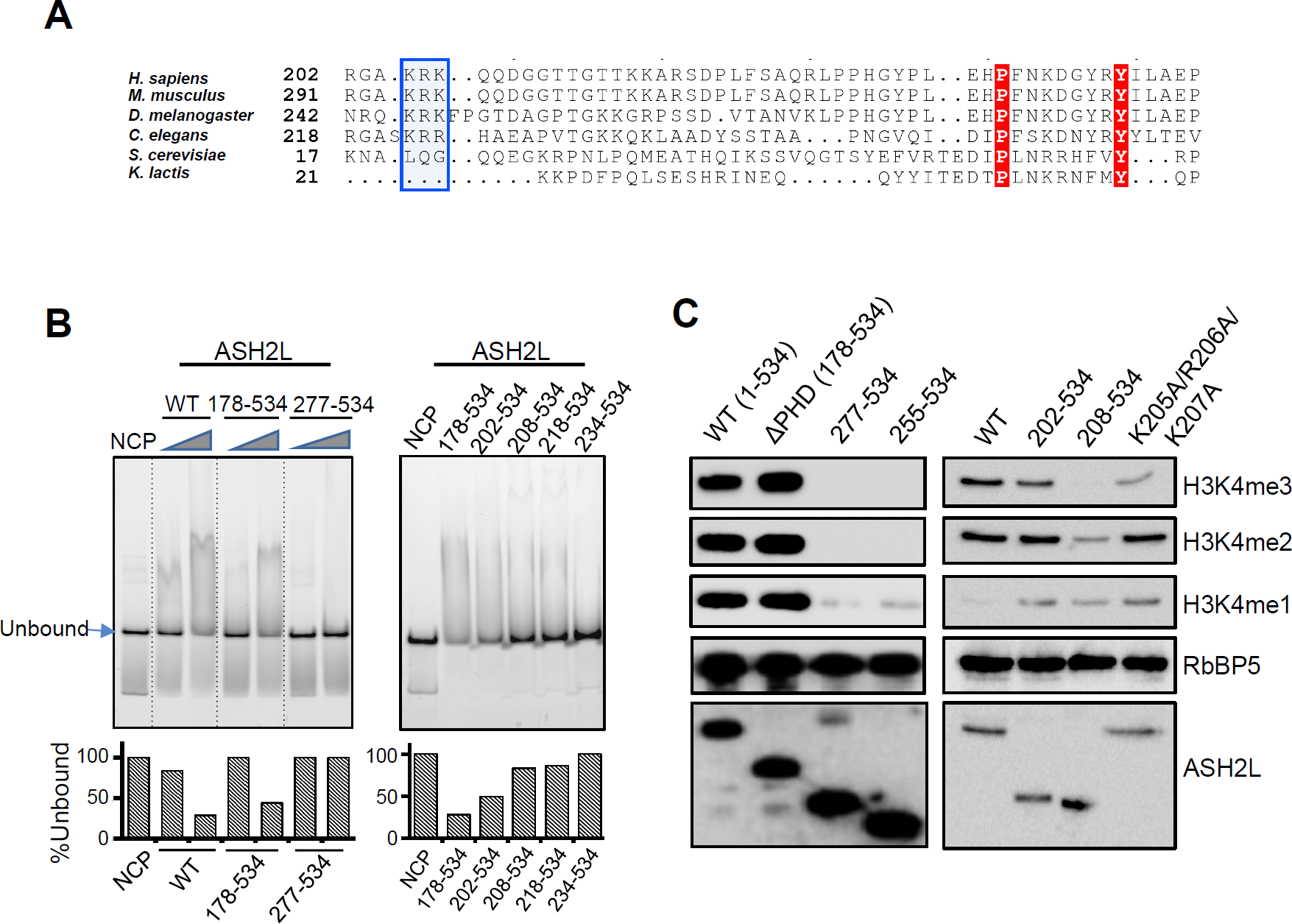
ASH2L Linker-IDR is Important for NCP Binding and Methyltransferase Activity. (A) Multiple sequence alignment of ASH2L Linker-IDR region (residues 202–254). The blue box indicated _205_-KRK-_207_, key residues for NCP recognition. (B) Electrophoretic mobility shift assay of ASH2L and ASH2L mutants as indicated on top. The unbound NCP quantification was done with ImageJ and normalized against the NCP lane on left, which was arbitrarily set as 1 (100%). (C) Immunoblot to detect *in vitro* histone methyltransferase activity with the NCP as the substrate. Reconstituted MLL1^RWSAD^ complexes containing wild type and mutant ASH2L, used as indicated on top. Immunoblots of RbBP5 and ASH2L included as controls.

### Alignment of MLL1^SET^ at the Nucleosome Dyad

Given the binding of RbBP5 and ASH2L at the edge of the NCP (SHL1.5 and SHL7), the catalytic MLL1^SET^ domain was positioned at the nucleosome dyad (Figure 6A). In this arrangement, the active site of the MLL1^SET^ domain pointed outward (Figure 6A). The NCP structure was well-resolved in the cryo-EM map of MLL1^RWSAD^-NCP. Both histone H3 tails emanated from between two gyres of nucleosomal DNA, with Lys37 as the first observable residue on histone H3 N-terminal tails (Figure 6A). The distance between Lys37 on each histone H3 tail and the active site of MLL1^SET^ was ∼ 60 Å. Thus, K4 residues on both H3 tails were almost equally accessible to the MLL1^SET^ active site (Figure 6A). The distance constraint restricted access of the MLL1^SET^ domain to only K4 and K9 on H3 N-terminus (Figure 6A). More importantly, since MLL1^SET^ is a non-processive enzyme (Patel et al., 2009), close proximity of the MLL1^SET^ domain to K4 on both H3 tails likely played a significant role in promoting its activity on higher H3K4me states on the NCP (Figure 6B, see discussion).

**Figure 6.**
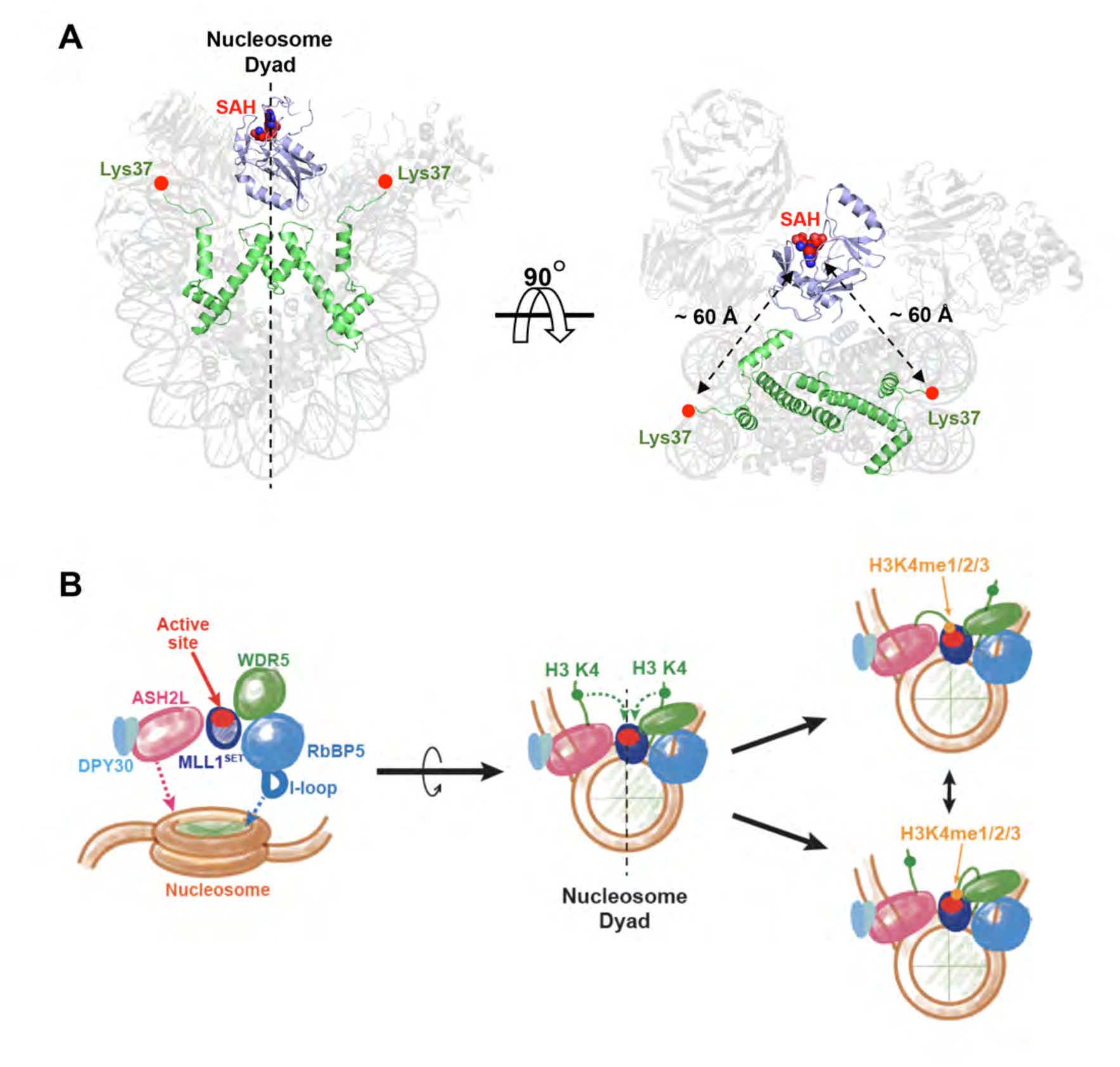
The MLL1^SET^ Domain Aligns at the Nucleosomal Dyad. (A) MLL1^SET^ (slate) and two copies of histone H3 (green) were highlighted against the faded MLL1^RWSAD^-NCP structure. The SAH molecule represented as a sphere and marked the catalytic active site of MLL1^SET^. Both histone H3 tails emanated from between two gyres of nucleosomal DNA (Lys37) maintained similar distance from the MLL1^SET^ active site (∼ 60 Å). (B) Schematic model of NCP recognition mediated by the MLL1 core complex. The I-loop of RbBP5 and the active site of MLL1^SET^ were colored in blue and red, respectively. NCP recognition by the MLL1 core complex functions mainly through concurrent DNA contacts via RbBP5 and ASH2L. The I-loop of RbBP5 and H4 tail interaction provided the specific docking point for MLL1 binding to the NCP. This allowed the active site of MLL1^SET^ to align at the nucleosome dyad. As a result, both histone H3 tails had physical proximity to MLL1^SET^, a non-processive enzyme, for successive rounds of methylation.

## DISCUSSION

Here we report the single-particle cryo-EM structure of the human MLL1 core complex bound to the NCP. It shows that the MLL1 core complex anchors on the NCP through WD40 domain of RbBP5 and Linker-IDR of ASH2L. This dual interaction positions the catalytic MLL1^SET^ domain near the nucleosome dyad, allowing symmetric access to both H3K4 substrates within the NCP. Disruption of MLL1-NCP interaction specifically reduces MLL1 activity for nucleosomal H3K4me3. Our study sheds light on how the MLL1 core complex engages chromatin and how chromatin binding promotes MLL1 tri-methylation activity.

### Unique MLL1-NCP Recognition among Chromatin Recognition Complexes

One of the well-known features of the NCP is the acidic patch, which is a negatively-charged and solvent exposed surface. It is organized by a series of negatively-charged residues in histones H2A and H2B (Zhou et al., 2019). The acidic patch interacts with the basic patch on histone H4 of adjacent nucleosomes, which underlies inter-nucleosome interactions in higher order chromatin structures (Kalashnikova et al., 2013). Structures of NCP-protein complexes demonstrate that the acidic patch is recognized by NCP-interacting proteins in many cases through diverse arginine finger motifs (e.g., LANA (Barbera et al., 2006), RCC1 (Makde et al., 2010), 53BP1 (Wilson et al., 2016)). Our study demonstrates that the MLL1 core complex binds to a unique surface of the NCP that do not involve the acidic patch. The main contributors to the NCP interaction are the electrostatic interactions between positively-charged residues in RbBP5 and ASH2L and the DNA backbone in the NCP. Extensive DNA-interactions were also observed for the histone H3K27 methyltransferase PRC2 in complex with dinucleosomes (Poepsel et al., 2018). However, unlike PRC2, all MLL1-NCP interactions occur within a single nucleosome. It is possible that other domains of MLL1 (e.g. MLL1 PHD-Bromo) that are not included in our study are important for engaging adjacent nucleosomes and spreading the H3K4me marks.

In our structure, the I-loop of RbBP5, inserting between the H4 tail and nucleosomal DNA (SHL1.5), provides the specific docking point of the MLL1 core complex to the NCP. Once RbBP5 is docked, the distance between RbBP5 and ASH2L (∼ 70 Å) limits ASH2L binding to SHL7 of the NCP (Figure S7C). Interestingly, despite importance of RbBP5 I-loop in specifying the orientation of MLL1 on the NCP, it did not contribute significantly to binding of RbBP5 to the NCP (Figure 2G). Nonetheless, dual recognition through both specific and non-specific interactions of RbBP5 and ASH2L likely enables the MLL1 core complex to bind the NCP in a unique configuration for optimal access to both H3K4 substrates (Figure 6B). Notably, H4 interactions are also used by other chromatin interacting proteins e.g., DOT1L, SNF, ISWI (Anderson et al., 2019; Jang et al., 2019; Li et al., 2019a; Valencia-Sanchez et al., 2019; Worden et al., 2019; Yan et al., 2019; Yao et al., 2019), raising the possibility of another NCP docking site in addition to the acidic patch.

### Structural Conservation and Divergence of the Human MLL1 Core Complex

The composition of catalytically-active human MLL/SET and yeast SET1 complexes is largely conserved (Rao and Dou, 2015). Our study shows that human MLL1 core complex and the yeast SET1 complexes (Hsu et al., 2018; Qu et al., 2018) have the same overall architecture, with the catalytic MLL1^SET^ domain sandwiched by RbBP5-WDR5 (Swd1-Swd3 in *Kluveromyces lactis*/Cps50-Cps30 in *Saccharomyces cerevisiae*) and ASH2L-DPY30 (Bre2-Sdc1/Cps60-Cps25 in yeast) on each side (Figure 3C). Furthermore, the crystal structure of the yeast SET1 complex (Hsu et al., 2018) overlays well with the MLL1 core complex in our structure. This may suggest that the yeast SET1 complex adopts a similar configuration on the NCP.

Given that RbBP5 and ASH2L are shared among all MLL family protein complexes in mammals, it is likely that our study reveals a general mechanism for how the mammalian MLL complexes engage chromatin and gain access to the H3K4 substrates. Sequence alignments show significant conservation of RbBP5 I-/A-loops as well as the basic residues in ASH2L in higher eukaryotes. It supports the functional importance of these regions in chromatin binding and H3K4me regulation. It also suggests that the mechanism by which MLL family enzymes engage chromatin in higher eukaryotes is likely to be conserved. Importantly, recent genome sequencing studies have identified mutations in these conserved regions in human malignancies (Blankin et al., 2012; Kim et al., 2014), which warrants future studies. We would like to point out that the interface of RbBP5 and ASH2L with the NCP are not well conserved in homologous yeast Swd1/Cps50 and Bre2/Cps60 protein (Figures 5A and S5B). The I-loop is much shorter and the A-loop is missing in the yeast Swd1/Cps50 protein (Figure S5A), suggesting potential divergence of detailed yeast SET1-NCP interactions at the molecular level, although the overall NCP recognition pattern might be similar.

### Contribution of ASH2L-IDR in NCP Recognition and H3K4me Regulation

Previous studies have shown that several components of the MLL1 core complex are capable of interacting with DNA or RNA, including WDR5, RbBP5 and ASH2L (Chen et al., 2011; Mittal et al., 2018; Sarvan et al., 2011; Yang et al., 2014). This raises the question of how these interactions contribute to recruitment of the MLL1 core complex to chromatin. Our structure indicates that WDR5, MLL1^SET^ as well as PHD-WH domain of ASH2L do not directly interact with the NCP, suggesting that these proteins probably interact with either nucleosome-free DNAs and indirectly contribute to the stability of the MLL1 core complex on chromatin. Our study reveals a previously uncharacterized function of ASH2L Linker-IDRs in chromatin function. This region was not studied in the previous MLL1^SET^-ASH2L^SPRY^-RbBP5^330-375^ structure (Li et al., 2016). The ASH2L IDRs contain evolutionarily conserved sequences (Figure 5A and S5B) and exhibit dynamic properties on the NCP. Notably, we take advantage of protein structure prediction approach to identify the essential interface between ASH2L IDRs and the NCP. This approach subsequently allowed us to uncover a basic patch region in ASH2L (_205_-KRK-_207_) that significantly contributes to MLL1 binding to the NCP and MLL1 tri-methylation activity. However, it is likely that other regions of ASH2L also contribute to NCP binding since deletion of the ASH2L residues 202–254 leads to more prominent reduction of H3K4me (Figure 5B-D). Molecular dissection of ASH2L IDRs awaits future studies.

### Regulation of MLL1 Activity for H3K4me3 on the NCP

Single turnover kinetic experiments revealed that the MLL1 core complex uses a non-processive mechanism for catalysis (Patel et al., 2009), requiring capture and release H3K4 after each round of the methylation reaction. Thus, it is not kinetically favorable to achieve tri-methylation state when enzyme and substrate have random encounters in solution. Our study here demonstrates that the MLL1 core complex stably associates with NCP via RbBP5 and ASH2L, which uniquely positions the MLL1^SET^ domain at the nucleosome dyad with near symmetric access to both H3K4 substrates (Figure 6A). Stable settlement on the NCP allows close physical proximity and optimal orientation of the MLL1^SET^ catalytic site to both H3K4 substrates on the NCP, which significantly favor the kinetics of successive methylation reactions. In support, disruption of the MLL1-NCP interactions significantly reduces H3K4me3 activity on the NCP without affecting MLL1 activity on free H3. Given enhanced MLL1 activity on the NCP, we envision that stabilizing the MLL1 complex on chromatin by transcription factors and cofactors will further enhances overall H3K4me3, which in turn recruits additional transcription cofactors (Lauberth et al., 2013; Ruthenburg et al., 2007; Taverna et al., 2007; Vermeulen et al., 2007; Wysocka et al., 2006). This leads to a feedback loop for optimal gene expression and spreading of H3K4me3 at the actively transcribed genes in cells. Position of the MLL1^SET^ domain at the nucleosome dyad also raises the question of potential interplay with linker histones, which bind near the nucleosome dyad in the heterochromatic regions in eukaryotes (Hayes et al., 1994; Lu et al., 2013; Zhou et al., 2015). It would be interesting to test whether linker histone inhibits MLL1 activity and thus promotes closed chromatin conformation in future.

## AUTHOR CONTRIBUTIONS

S.H.P and A.A. designed, performed experiments and wrote the manuscript. A.A. Y.T.L. and J.X performed *in vitro* biochemical experiments and contributed to experimental design; W.Z., B.Z. and Y.Z performed molecular simulation analyses; A.A., H.K. and S.A. prepared proteins and helped with cryo-EM samples. M.A.C. and M.S. contributed to cryo-EM data acquisition and analysis; Y.D and U.S.C conceptualized and supervised the overall study and wrote the manuscript.

## ACKNOWLEDGMENTS

We thank Dr. Amy Bondy at University of Michigan cryo-EM center for assistance with data collection. We thank Dr. Takanori Nakane and Dr. Jeong Min Chung for helping with cryo-EM data processing. This work was supported by grants (GM082856) to Y.D. and (DK111465 and NRF-2015M3D3A1A01064876) to U.S.C. The authors declare no competing financial interests.

## EXPERIMENTAL PROCEDURES

### Protein expression and purification

The core subunits of the MLL1^RWSAD^ complex (MLL1^SET,3762-3969^ ASH2L^1-534^ and mutants, RbBP5^1-538^ and mutants, WDR5^23-334^, and DPY30^1-99^) were expressed and purified using the pET28a His_6_-small ubiquitin-related modifier (SUMO) vector as previously described (Cao et al., 2010; Li et al., 2016). Mutations of RbBP5 and ASH2L were generated by overlap PCR-based site-directed mutagenesis. Individual components of the MLL1^RWSAD^ complex were purified on Ni-NTA column and equimolar quantities were mixed and purified by gel filtration chromatography as previously described (Cao et al., 2010; Li et al., 2016). Full-length *X. laevis* histones H2A, H2B, H3, and H4 were expressed and purified using the one-pot purification method (Lee et al., 2015). Assembly of the nucleosome core particle using 147 base-pair Widom 601 DNA (Lowary and Widom, 1998) was done by salt dialysis as previously described (Luger et al., 1997; Luger et al., 1999).

### *In vitro* histone methyltransferase assay

The *in vitro* histone methyltransferase assay was carried out by incubating the MLL1^RWSAD^ complex (0.3 µM) with either nucleosome (0.965 µM) or free recombinant histone H3 (0.098 µM) for 1 hour at room temperature. The reaction buffer contained 20 mM Tris-HCl, pH 8.0, 50 mM NaCl, 1 mM DTT, 5 mM MgCl_2_ and 10% v/v glycerol in a total volume of 20 µL. Reactions were quenched with 20 µL of 2X Laemmli Sample Buffer (Bio-Rad cat. #161-0737). H3K4 methylation was detected by western blot using antibodies for H3K4me_1_ (1:20000, Abcam cat. No. ab8895), H3K4me_2_ (1:40000, EMD-Millipore cat.#07-030), or H3K4me_3_ (1:10000, EMD-Millipore cat.#07-473) for either 1hr at room temperature or overnight at 4 °C. The blot was then incubated with IgG-HRP (Santa Cruz Biotechnology cat.#sc-486) for 1 h at room temperature. The membrane was developed using ECL (Pierce cat.#32106) and visualized by chemiluminescence (Bio-Rad ChemiDoc Imaging System).

### Electrophoretic Mobility Shift Assay (EMSA)

EMSA assay was carried out using 0.1 µM nucleosomes and increasing concentration of MLL1 subunits. The protein mixture was run on the 6% 0.2X TBE gel that was pre-run for 1.5 hours, 150 V at 4 °C. The gel was visualized by incubating in 100 mL of TAE with 1:20000 diluted ethidium bromide for 10 minutes at room temperature. Gels were then incubated in distilled water for 10 minutes and visualized by UV transillumination (Bio-Rad ChemiDoc Imaging System). The results were quantified by ImageJ software (Schneider et al., 2012).

### His_6_ Pull-down Assay

His_6_-fusion proteins were incubated with the NCP in BC150 (20 mM Tris-HCl, pH 7.5, 350 mM NaCl, 20 mM imidazole, 0.05% v/v NP-40, 10 mM DTT, 1 mg/ml BSA, PMSF and inhibitor cocktail) for 2 hours at 4 °C. After several washes with BC150, the beads were boiled in SDS loading buffer and analyzed by Western blot.

### Cryo-EM sample preparation

The cryo-EM sample was prepared by the GraFix method (Kastner et al., 2008). Specifically, the reconstituted MLL1^RWSAD^ complex (30 µM) was incubated with nucleosomes (10 µM) in GraFix buffer (50 mM HEPES, pH7.5, 50 mM NaCl, 1 mM MgCl_2_, 1 mM TECP) with added 0.5 mM SAH for 30 min at 4 °C. The sample was applied onto the top of the gradient solution (0-60% glycerol gradient with 0-0.2% glutaraldehyde, in GraFix buffer) and was centrifuged at 48,000 rpm at 4 °C for 3 hours. After ultracentrifugation, 20 µl fractions were manually collected from the top of the gradient. The crosslinking reaction was terminated by adding 2 µl of 1 M Tris-HCl, pH7.5 into each fraction. Glycerol was removed by dialyzing the sample in GraFix buffer using centrifugal concentrator (Sartorius Vivaspin 500) before making cryo-EM grids.

### Cryo-EM data collection and processing

A protein sample at 1 mg/ml concentration was plunge-frozen on 200 mesh quantifoil R1.2/1.3 grids (Electron Microscopy Sciences) using a Mark IV Vitrobot (Thermo Fisher Scientific) with settings as 4 °C, 100% humidity and 4 sec blotting time. Cryo-EM grids were imaged on a FEI Titan Krios operating at 300 KV at liquid nitrogen temperature. The Gatan K2 Summit direct electron detector was used at a nominal magnification of 29,000 X in a counting mode with a pixel size of 1.01 Å/pixel. A dose rate of 8 electrons/Å^2^/s and defocus values ranging from −1.5 to −3.5 µm were used. Total exposure of 8 sec per image was dose-fractionated into 40 movie frames, resulting in an accumulated dose of 64 electrons per Å^2^. A total of 4717 movies were collected for the MLL1^RWSAD^-NCP dataset.

Micrograph movie stacks were first subjected to MotionCor2 for whole-frame and local drift correction (Zheng et al., 2017). For each micrograph, CTFFIND4.1 was used to fit the contrast transfer function (Rohou and Grigorieff, 2015). The estimated resolution of micrographs lower than 5Å were excluded from further processing, which resulted in 3896 micrographs. Particle picking was performed using the Warp (Tegunov and Cramer, 2018), which picked total 712,198 particles. Using particle coordinates obtained from the Warp, the particles were extracted with the box size of 350 Å using RELION 3 program package (Zivanov et al., 2018). Extracted particles were then imported into cryoSPARC (Punjani et al., 2017) for 2D classification in 200 classes. After removal of bad classes, the total of 694,180 particles were subjected to *ab initio* 3D classification (Figure S2). The major class (323,408 particles) contained the MLL1 core complex and the NCP, which was then subjected for the heterogeneous refinement. This led to the identification of ten subclasses. One subclass showed the partial cryo-EM density for the MLL1 core complex, thus excluded for the further processing. The remaining nine subclasses (252,109 particles) maintained intact MLL1^RWSAD^-NCP complex. These nine subclasses also used for the rigid-body fitting of individual component of the MLL1^RWSAD^-NCP complex to visualize the dynamics of each component against the NCP (Figure S3E). 252,109 particles were imported in RELION and performed the 3D classification without alignment (10 classes, 35 cycles, T=40). One out of 10 classes (8,433 particles) exhibited the well-defined map of MLL1^RWSAD^-NCP. These particles were used for 3D refinement in RELION and post-processed to a resolution of 6.2 Å and a B factor of −189 Å^2^. This cryo-EM map was local filtered using RELION to the local resolution to avoid over-interpretation.

To obtain a cryo-EM map for RbBP5-NCP and MLL1^RWS^-NCP subcomplexes, we utilized RELION’s multi-body refinement procedure with 252,109 particles (Figure S2). RbBP5-NCP (32,563 particles), MLL1^WSAD^, MLL1^RWS^-NCP (21,114 particles), and MLL1^AD^ were separately masked during the multibody refinement (Nakane et al., 2018). The partial signal subtraction was performed to generate the particle set for RbBP5-NCP and MLL1^RWS^-NCP. Further 3D classifications without alignment (5 classes, 35 cycles, T=40 for RbBP5-NCP and 10 classes, 35 cycles, T=40 for MLL1^RWS^-NCP) were performed and the best maps based on the resolution and occupancy of RbBP5 and MLL1^RWS^ densities were selected for further refinement and post-processing (Figure S2). The reported final resolution of each cryo-EM structure was estimated by RELION with Fourier shell correlation (FSC) at criteria of 0.143 (Figure S3A-C).

### Modeling, rigid body fitting, and model refinement

We built a 3D atomic model of the human ASH2L protein by I-TASSER (Roy et al., 2010; Wei Zheng, 2019; Zhang, 2008) assisted by deep-learning based contact-map prediction (Li et al., 2019b). The fragment-guided molecular dynamics refinement software, FG-MD (Zhang et al., 2011), was utilized to remove the steric clash between ASH2L model and other molecules and further refine the local structures (Figure S6B). Finally, our in-house EM-fitting software, EM-Ref (Zhang et al, in preparation), was used to fit the ASH2L model and other parts of human MLL1 core complex to the density maps to get final atomic models.

I-TASSER utilized LOMETS, which consisted of 16 individual threading programs (Zheng et al., 2019), to generate templates as the initial conformation. Human ASH2L protein consisted of three domains, while the 2^nd^, 3^rd^ domains (Linker-IDR and ASH2L^SPRY^) and C-terminal SDI motif can be covered by templates (PDB ID: 6E2H and 6CHG, B chain, crystal structure of the yeast SET1 H3K4 methyltransferase catalytic module (Hsu et al., 2018)) in most of the top threading alignments. The 1^st^ domain (PHD-WH domain) was covered by another template (PDB ID: 3S32, A chain, the crystal structure of ASH2L N-terminal domain) (Sarvan et al., 2011). Therefore, these three proteins were used as the main templates for building the full-length ASH2L model, where structural assembly simulation was guided by the contact-maps from the deep-learning program, ResPRE (Li et al., 2019). Finally, the first model of I-TASSER was selected as the potential ASH2L model, where the estimated TM-score (Xu and Zhang, 2010) for the C-terminal domain is 0.71±0.12, suggesting that the confidence of the I-TASSER model is high. Superposing ASH2L model (Linker-IDR and ASH2L^SPRY^) with the experimental structure (ASH2L^SPRY^) is shown in Figure 4A.

Monte Carlo (MC) simulation was employed to fit and refine the complex model structures based on the experimental density map. During the MC simulations, individual domain structures were kept as the rigid-body, where global translation and rotation of the domains were performed, which would be accepted or rejected based on Metropolis algorithm (Binder et al., 1993). The total number of translation and rotation was 50,000 in the MC simulation. The MC energy function used in the simulation was a linear combination of correlation coefficient (CC) between structural models and the density map data and the steric clashes between the atomic structures, i.e.,

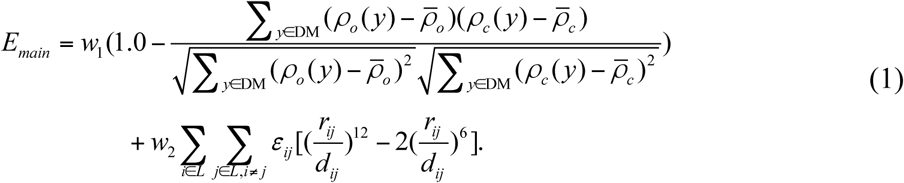

Where ρ_c_(y) was the calculated density map on grid (DiMaio et al., 2009). ρ_o_(y) was obtained from the experimental density map. 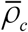 and 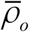 were the average of calculated density map and experimental density map, respectively. *DM* and *L* represented tbe density map and the length of protein, respectively. *d*_*ij*_ was the distance between the two atoms *i* and *j. r*_*ij*_ was the sum of their van der Waals atomic radii and *ε*_*ij*_ was the combined well-depth parameter for atoms *i* and *j*, which were all taken from the CHARMM force field (MacKerell Jr et al., 1998). *w*_*1*_=100 and *w*_*2*_=1 were the weights for correlation coefficient item and clash item, respectively.

For the nucleosome model, the crystal structure of nucleosome (PDB ID:3MVD) (Makde et al, 2010) was used for rigid-body fitting. In the cryo-EM structure of RbBP5-NCP, the histone H4 tail region was manually rebuilt where the density allowed using the program COOT (Emsley et al., 2010). Three model structures of MLL1^RWSAD^-NCP, RbBP5-NCP, and MLL1^RWS^-NCP were subjected to the real-space refinement using PHENIX after rigid-body fitting. Validations of three model structures were performed by MolProbity (Chen et al., 2010). The final structures were further validated by calculating map-model FSC curves using phenix.mtriage in the PHENIX program package (Figure S3D) (Afonine et al., 2018). The computed FSC between the model and map agreed reasonably well as shown in Figure S3D. Statistics for data collection, refinement, and validation summarized in Table 1.

### DATA Accessions

The accession numbers for the MLL1^RWSAD^-NCP, RbBP5-NCP and MLL1^RWS^-NCP cryo-EM structures are PDB: 6PWV and EMDB: EMD-20512; PDB 6PWX and EMDB: EMD-20514; and PDB: 6PWW and EMDB: EMD-20513, respectively.

## REFERENCES

Adams, P.D., Afonine, P.V., Bunko′ czi, G., Chen, V.B., Davis, I.W., Echols, N., Headd, J.J., Hung, L.W., Kapral, G.J., Grosse-Kunstleve, R.W., et al. (2010). PHENIX: a comprehensive Python-based system for macromolecular structure solution. Acta Crystallogr. D Biol. Crystallogr. 66, 213–221.

Afonine, P.V., Klaholz, B.P., Moriarty, N.W., Poon, B.K., Sobolev, O.V., Terwilliger, T.C., Adams, P.D., and Urzhumtsev, A. (2018). New tools for the analysis and validation of cryo-EM maps and atomic models. Acta Crystallogr. D Struct. Biol. 74, 814–840.

Anderson, C.J., Baird, M.R., Hsu, A., Barbour, E.H., Koyama, Y., Borgnia, M.J., and McGinty, R.K. (2019). Structural Basis for Recognition of Ubiquitylated Nucleosome by Dot1L Methyltransferase. Cell Rep. 26, 1681–1690 e1685.

Armache, K.J., Garlick, J.D., Canzio, D., Narlikar, G.J., and Kingston, R.E. (2011). Structural basis of silencing: Sir3 BAH domain in complex with a nucleosome at 3.0 A resolution. Science 334, 977–982.

Avdic, V., Zhang, P., Lanouette, S., Groulx, A., Tremblay, V., Brunzelle, J., and Couture, J.F. (2011). Structural and biochemical insights into MLL1 core complex assembly. Structure 19, 101–108.

Barbera, A.J., Chodaparambil, J.V., Kelley-Clarke, B., Joukov, V., Walter, J.C., Luger, K., and Kaye, K.M. (2006). The nucleosomal surface as a docking station for Kaposi’s sarcoma herpesvirus LANA. Science 311, 856–861.

Binder, K., Heermann, D., Roelofs, L., Mallinckrodt, A.J., and McKay, S. (1993). Monte Carlo simulation in statistical physics. Computers in Physics 7, 156–157.

Bochynska, A., Luscher-Firzlaff, J., and Luscher, B. (2018). Modes of Interaction of KMT2 Histone H3 Lysine 4 Methyltransferase/COMPASS Complexes with Chromatin. Cells 7. doi: 10.3390/cells7030017.

Butler, J.S., Qiu, Y.H., Zhang, N., Yoo, S.Y., Coombes, K.R., Dent, S.Y., and Kornblau, S.M. (2017). Low expression of ASH2L protein correlates with a favorable outcome in acute myeloid leukemia. Leuk. Lymphoma 58, 1207–1218.

Cao, F., Chen, Y., Cierpicki, T., Liu, Y., Basrur, V., Lei, M., and Dou, Y. (2010). An Ash2L/RbBP5 heterodimer stimulates the MLL1 methyltransferase activity through coordinated substrate interactions with the MLL1 SET domain. PLoS One 5, e14102.

Chen, K., Chen, Z., Wu, D., Zhang, L., Lin, X., Su, J., Rodriguez, B., Xi, Y., Xia, Z., Chen, X., et al. (2015). Broad H3K4me3 is associated with increased transcription elongation and enhancer activity at tumor-suppressor genes. Nat. Genet. 47, 1149–1157.

Chen, V.B., Arendall, W.B., 3rd, Headd, J.J., Keedy, D.A., Immormino, R.M., Kapral, G.J., Murray, L.W., Richardson, J.S., and Richardson, D.C. (2010). MolProbity: all-atom structure validation for macromolecular crystallography. Acta Crystallogr. D Biol. Crystallogr. 66, 12–21.

Chen, Y., Wan, B., Wang, K.C., Cao, F., Yang, Y., Protacio, A., Dou, Y., Chang, H.Y., and Lei, M. (2011). Crystal structure of the N-terminal region of human Ash2L shows a winged-helix motif involved in DNA binding. EMBO Rep. 12, 797–803.

Cosgrove, M.S., and Patel, A. (2010). Mixed lineage leukemia: a structure-function perspective of the MLL1 protein. FEBS J. 277, 1832–1842.

Couture, J.F., Collazo, E., and Trievel, R.C. (2006). Molecular recognition of histone H3 by the WD40 protein WDR5. Nat. Struct. Mol. Biol. 13, 698–703.

DiMaio, F., Tyka, M.D., Baker, M.L., Chiu, W., and Baker, D. (2009). Refinement of protein structures into low-resolution density maps using rosetta. J. Mol. Biol. 392, 181–190.

Dou, Y., Milne, T.A., Tackett, A.J., Smith, E.R., Fukuda, A., Wysocka, J., Allis, C.D., Chait, B.T., Hess, J.L., and Roeder, R.G. (2005). Physical association and coordinate function of the H3 K4 methyltransferase MLL1 and the H4 K16 acetyltransferase MOF. Cell 121, 873–885.

Emsley, P., Lohkamp, B., Scott, W.G., and Cowtan, K. (2010). Features and development of Coot. Acta Crystallogr. D Biol. Crystallogr. 66, 486–501.

Ge, Z., Song, E.J., Kawasawa, Y.I., Li, J., Dovat, S., and Song, C. (2016). WDR5 high expression and its effect on tumorigenesis in leukemia. Oncotarget 7, 37740–37754.

Goddard, T.D., Huang, C.C., Meng, E.C., Pettersen, E.F., Couch, G.S., Morris, J.H., and Ferrin, T.E. (2018). UCSF ChimeraX: Meeting modern challenges in visualization and analysis. Protein Sci. 27, 14–25.

Guenther, M.G., Jenner, R.G., Chevalier, B., Nakamura, T., Croce, C.M., Canaani, E., and Young, R.A. (2005). Global and Hox-specific roles for the MLL1 methyltransferase. Proc. Natl. Acad. Sci. USA 102, 8603–8608.

Haddad, J.F., Yang, Y., Takahashi, Y.H., Joshi, M., Chaudhary, N., Woodfin, A.R., Benyoucef, A., Yeung, S., Brunzelle, J.S., Skiniotis, G., et al. (2018). Structural Analysis of the Ash2L/Dpy-30 Complex Reveals a Heterogeneity in H3K4 Methylation. Structure 26, 1594–1603 e1594.

Hannibal, M.C., Buckingham, K.J., Ng, S.B., Ming, J.E., Beck, A.E., McMillin, M.J., Gildersleeve, H.I., Bigham, A.W., Tabor, H.K., Mefford, H.C., et al. (2011). Spectrum of MLL2 (ALR) mutations in 110 cases of Kabuki syndrome. Am J Med Genet A 155A, 1511–1516.

Hayes, J.J., Pruss, D., and Wolffe, A.P. (1994). Contacts of the globular domain of histone H5 and core histones with DNA in a “chromatosome”. Proc. Natl. Acad. Sci. USA 91, 7817–7821.

Hsu, P.L., Li, H., Lau, H.T., Leonen, C., Dhall, A., Ong, S.E., Chatterjee, C., and Zheng, N. (2018). Crystal Structure of the COMPASS H3K4 Methyltransferase Catalytic Module. Cell 174, 1106–1116 e1109.

Jang, S., Kang, C., Yang, H.S., Jung, T., Hebert, H., Chung, K.Y., Kim, S.J., Hohng, S., and Song, J.J. (2019). Structural basis of recognition and destabilization of the histone H2B ubiquitinated nucleosome by the DOT1L histone H3 Lys79 methyltransferase. Genes Dev. 33, 620–625.

Jones, W.D., Dafou, D., McEntagart, M., Woollard, W.J., Elmslie, F.V., Holder-Espinasse, M., Irving, M., Saggar, A.K., Smithson, S., Trembath, R.C., et al. (2012). De novo mutations in MLL cause Wiedemann-Steiner syndrome. Am. J. Hum. Genet. 91, 358–364.

Kalashnikova, A.A., Porter-Goff, M.E., Muthurajan, U.M., Luger, K., and Hansen, J.C. (2013). The role of the nucleosome acidic patch in modulating higher order chromatin structure. J. R. Soc. Interface 10, 20121022.

Kastner, B., Fischer, N., Golas, M.M., Sander, B., Dube, P., Boehringer, D., Hartmuth, K., Deckert, J., Hauer, F., Wolf, E., et al. (2008). GraFix: sample preparation for single-particle electron cryomicroscopy. Nat. Methods 5, 53–55.

Katada, S., and Sassone-Corsi, P. (2010). The histone methyltransferase MLL1 permits the oscillation of circadian gene expression. Nat. Struct. Mol. Biol. 17, 1414–1421.

Kato, H., Jiang, J., Zhou, B.R., Rozendaal, M., Feng, H., Ghirlando, R., Xiao, T.S., Straight, A.F., and Bai, Y. (2013). A conserved mechanism for centromeric nucleosome recognition by centromere protein CENP-C. Science 340, 1110–1113.

Kluijt, I., van Dorp, D.B., Kwee, M.L., Toutain, A., Keppler-Noreuil, K., Warburg, M., and Bitoun, P. (2000). Kabuki syndrome - report of six cases and review of the literature with emphasis on ocular features. Ophthalmic Genet. 21, 51–61.

Kornberg, R.D., and Thomas, J.O. (1974). Chromatin structure; oligomers of the histones. Science 184, 865–868.

Krajewski, W.A., Li, J., and Dou, Y. (2018). Effects of histone H2B ubiquitylation on the nucleosome structure and dynamics. Nucleic Acids Res. 46, 7631–7642.

Lauberth, S.M., Nakayama, T., Wu, X., Ferris, A.L., Tang, Z., Hughes, S.H., and Roeder, R.G. (2013). H3K4me3 interactions with TAF3 regulate preinitiation complex assembly and selective gene activation. Cell 152, 1021–1036.

Lee, Y.T., Gibbons, G., Lee, S.Y., Nikolovska-Coleska, Z., and Dou, Y. (2015). One-pot refolding of core histones from bacterial inclusion bodies allows rapid reconstitution of histone octamer. Protein Expr. Purif. 110, 89–94.

Li, M., Xia, X., Tian, Y., Jia, Q., Liu, X., Lu, Y., Li, M., Li, X., and Chen, Z. (2019a). Mechanism of DNA translocation underlying chromatin remodelling by Snf2. Nature 567, 409–413.

Li, Y., Bogershausen, N., Alanay, Y., Simsek Kiper, P.O., Plume, N., Keupp, K., Pohl, E., Pawlik, B., Rachwalski, M., Milz, E., et al. (2011). A mutation screen in patients with Kabuki syndrome. Hum. Genet. 130, 715–724.

Li, Y., Han, J., Zhang, Y., Cao, F., Liu, Z., Li, S., Wu, J., Hu, C., Wang, Y., Shuai, J., et al. (2016). Structural basis for activity regulation of MLL family methyltransferases. Nature 530, 447–452.

Li, Y., Hu, J., Zhang, C., Yu, D.-J., and Zhang, Y. (2019b). ResPRE: high-accuracy protein contact prediction by coupling precision matrix with deep residual neural networks. Bioinformatics. doi: 10.1093/bioinformatics/btz291.

Lowary, P.T., and Widom, J. (1998). New DNA sequence rules for high affinity binding to histone octamer and sequence-directed nucleosome positioning. J. Mol. Biol. 276, 19–42.

Lu, X., Wontakal, S.N., Kavi, H., Kim, B.J., Guzzardo, P.M., Emelyanov, A.V., Xu, N., Hannon, G.J., Zavadil, J., Fyodorov, D.V., et al. (2013). Drosophila H1 regulates the genetic activity of heterochromatin by recruitment of Su(var)3-9. Science 340, 78–81.

Luger, K., Mader, A.W., Richmond, R.K., Sargent, D.F., and Richmond, T.J. (1997). Crystal structure of the nucleosome core particle at 2.8 A resolution. Nature 389, 251–260.

Luger, K., Rechsteiner, T.J., and Richmond, T.J. (1999). Preparation of nucleosome core particle from recombinant histones. Methods Enzymol. 304, 3–19.

MacKerell Jr, A.D., Bashford, D., Bellott, M., Dunbrack Jr, R.L., Evanseck, J.D., Field, M.J., Fischer, S., Gao, J., Guo, H., and Ha, S. (1998). All-atom empirical potential for molecular modeling and dynamics studies of proteins. J. Phys. Chem. B 102, 3586–3616.

Magerl, C., Ellinger, J., Braunschweig, T., Kremmer, E., Koch, L.K., Holler, T., Buttner, R., Luscher, B., and Gutgemann, I. (2010). H3K4 dimethylation in hepatocellular carcinoma is rare compared with other hepatobiliary and gastrointestinal carcinomas and correlates with expression of the methylase Ash2 and the demethylase LSD1. Hum. Pathol. 41, 181–189.

Makde, R.D., England, J.R., Yennawar, H.P., and Tan, S. (2010). Structure of RCC1 chromatin factor bound to the nucleosome core particle. Nature 467, 562–566.

Mendelsohn, B.A., Pronold, M., Long, R., Smaoui, N., and Slavotinek, A.M. (2014). Advanced bone age in a girl with Wiedemann-Steiner syndrome and an exonic deletion in KMT2A (MLL). Am. J. Med. Genet. A 164A, 2079–2083.

Micale, L., Augello, B., Fusco, C., Selicorni, A., Loviglio, M.N., Silengo, M.C., Reymond, A., Gumiero, B., Zucchetti, F., D’Addetta, E.V., et al. (2011). Mutation spectrum of MLL2 in a cohort of Kabuki syndrome patients. Orphanet. J. Rare Dis. 6, 38.

Mittal, A., Hobor, F., Zhang, Y., Martin, S.R., Gamblin, S.J., Ramos, A., and Wilson, J.R. (2018). The structure of the RbBP5 beta-propeller domain reveals a surface with potential nucleic acid binding sites. Nucleic Acids Res. 46, 3802–3812.

Nakane, T., Kimanius, D., Lindahl, E., and Scheres, S.H. (2018). Characterisation of molecular motions in cryo-EM single-particle data by multi-body refinement in RELION. Elife 7. doi: 10.7554/eLife.36861.

Ng, S.B., Bigham, A.W., Buckingham, K.J., Hannibal, M.C., McMillin, M.J., Gildersleeve, H.I., Beck, A.E., Tabor, H.K., Cooper, G.M., Mefford, H.C., et al. (2010). Exome sequencing identifies MLL2 mutations as a cause of Kabuki syndrome. Nat. genet. 42, 790–793.

Patel, A., Dharmarajan, V., Vought, V.E., and Cosgrove, M.S. (2009). On the mechanism of multiple lysine methylation by the human mixed lineage leukemia protein-1 (MLL1) core complex. J. Biol. Chem. 284, 24242–24256.

Paulussen, A.D., Stegmann, A.P., Blok, M.J., Tserpelis, D., Posma-Velter, C., Detisch, Y., Smeets, E.E., Wagemans, A., Schrander, J.J., van den Boogaard, M.J., et al. (2011). MLL2 mutation spectrum in 45 patients with Kabuki syndrome. Hum. Mutat. 32, E2018–2025.

Pettersen, E.F., Goddard, T.D., Huang, C.C., Couch, G.S., Greenblatt, D.M., Meng, E.C., and Ferrin, T.E. (2004). UCSF Chimera--a visualization system for exploratory research and analysis. J. Comput. Chem. 25, 1605–1612.

Poepsel, S., Kasinath, V., and Nogales, E. (2018). Cryo-EM structures of PRC2 simultaneously engaged with two functionally distinct nucleosomes. Nat. Struct. Mol. Biol. 25, 154–162.

Punjani, A., Rubinstein, J.L., Fleet, D.J., and Brubaker, M.A. (2017). cryoSPARC: algorithms for rapid unsupervised cryo-EM structure determination. Nat. Methods 14, 290–296.

Qu, Q., Takahashi, Y.H., Yang, Y., Hu, H., Zhang, Y., Brunzelle, J.S., Couture, J.F., Shilatifard, A., and Skiniotis, G. (2018). Structure and Conformational Dynamics of a COMPASS Histone H3K4 Methyltransferase Complex. Cell 174, 1117–1126 e1112.

Rao, R.C., and Dou, Y. (2015). Hijacked in cancer: the KMT2 (MLL) family of methyltransferases. Nat. Rev. Cancer 15, 334–346.

Rea, S., Eisenhaber, F., O’Carroll, D., Strahl, B.D., Sun, Z.W., Schmid, M., Opravil, S., Mechtler, K., Ponting, C.P., Allis, C.D., et al. (2000). Regulation of chromatin structure by site-specific histone H3 methyltransferases. Nature 406, 593–599.

Rohou, A., and Grigorieff, N. (2015). CTFFIND4: Fast and accurate defocus estimation from electron micrographs. J. Struct. Biol. 192, 216–221.

Roy, A., Kucukural, A., and Zhang, Y. (2010). I-TASSER: a unified platform for automated protein structure and function prediction. Nat. Protoc. 5, 725–738.

Ruthenburg, A.J., Allis, C.D., and Wysocka, J. (2007). Methylation of lysine 4 on histone H3: intricacy of writing and reading a single epigenetic mark. Mol. Cell 25, 15–30.

Sarvan, S., Avdic, V., Tremblay, V., Chaturvedi, C.P., Zhang, P., Lanouette, S., Blais, A., Brunzelle, J.S., Brand, M., and Couture, J.F. (2011). Crystal structure of the trithorax group protein ASH2L reveals a forkhead-like DNA binding domain. Nat. Struct. Mol. Biol. 18, 857–859.

Schneider, C.A., Rasband, W.S., and Eliceiri, K.W. (2012). NIH Image to ImageJ: 25 years of image analysis. Nat. Methods 9, 671–675.

Southall, S.M., Wong, P.S., Odho, Z., Roe, S.M., and Wilson, J.R. (2009). Structural basis for the requirement of additional factors for MLL1 SET domain activity and recognition of epigenetic marks. Mol. Cell 33, 181–191.

Strom, S.P., Lozano, R., Lee, H., Dorrani, N., Mann, J., O’Lague, P.F., Mans, N., Deignan, J.L., Vilain, E., Nelson, S.F., et al. (2014). De Novo variants in the KMT2A (MLL) gene causing atypical Wiedemann-Steiner syndrome in two unrelated individuals identified by clinical exome sequencing. BMC Med. Genet. 15, 49.

Taverna, S.D., Li, H., Ruthenburg, A.J., Allis, C.D., and Patel, D.J. (2007). How chromatin-binding modules interpret histone modifications: lessons from professional pocket pickers. Nat. Struct. Mol. Biol. 14, 1025–1040.

Tegunov, D., and Cramer, P. (2018). Real-time cryo-EM data pre-processing with Warp. BioRxiv 338558.

Valencia-Sanchez, M.I., De Ioannes, P., Wang, M., Vasilyev, N., Chen, R., Nudler, E., Armache, J.P., and Armache, K.J. (2019). Structural Basis of Dot1L Stimulation by Histone H2B Lysine 120 Ubiquitination. Mol. Cell 74, 1010–1019 e1016.

van Nuland, R., Smits, A.H., Pallaki, P., Jansen, P.W., Vermeulen, M., and Timmers, H.T. (2013). Quantitative dissection and stoichiometry determination of the human SET1/MLL histone methyltransferase complexes. Mol. Cell Biol. 33, 2067–2077.

Vermeulen, M., Mulder, K.W., Denissov, S., Pijnappel, W.W., van Schaik, F.M., Varier, R.A., Baltissen, M.P., Stunnenberg, H.G., Mann, M., and Timmers, H.T. (2007). Selective anchoring of TFIID to nucleosomes by trimethylation of histone H3 lysine 4. Cell 131, 58–69.

Zheng, W., Zhang, C., Pearce, R., Mortuza, S.M., and Zhang, Y. (2019). Deep-learning contact-map guided protein structure prediction in CASP13. PROTEINS: Structure, Function, and Bioinformatics, in press.

Wilson, M.D., Benlekbir, S., Fradet-Turcotte, A., Sherker, A., Julien, J.P., McEwan, A., Noordermeer, S.M., Sicheri, F., Rubinstein, J.L., and Durocher, D. (2016). The structural basis of modified nucleosome recognition by 53BP1. Nature 536, 100–103.

Worden, E.J., Hoffmann, N.A., Hicks, C.W., and Wolberger, C. (2019). Mechanism of Cross-talk between H2B Ubiquitination and H3 Methylation by Dot1L. Cell 176, 1490–1501 e1412.

Wysocka, J., Swigut, T., Milne, T.A., Dou, Y., Zhang, X., Burlingame, A.L., Roeder, R.G., Brivanlou, A.H., and Allis, C.D. (2005). WDR5 associates with histone H3 methylated at K4 and is essential for H3 K4 methylation and vertebrate development. Cell 121, 859–872.

Wysocka, J., Swigut, T., Xiao, H., Milne, T.A., Kwon, S.Y., Landry, J., Kauer, M., Tackett, A.J., Chait, B.T., Badenhorst, P., et al. (2006). A PHD finger of NURF couples histone H3 lysine 4 trimethylation with chromatin remodelling. Nature 442, 86–90.

Xu, J., and Zhang, Y. (2010). How significant is a protein structure similarity with TM-score = 0.5? Bioinformatics 26, 889–895.

Yan, L., Wu, H., Li, X., Gao, N., and Chen, Z. (2019). Structures of the ISWI-nucleosome complex reveal a conserved mechanism of chromatin remodeling. Nat. Struct. Mol. Biol. 26, 258–266.

Yang, Y.W., Flynn, R.A., Chen, Y., Qu, K., Wan, B., Wang, K.C., Lei, M., and Chang, H.Y. (2014). Essential role of lncRNA binding for WDR5 maintenance of active chromatin and embryonic stem cell pluripotency. Elife 3, e02046.

Yao, T., Jing, W., Hu, Z., Tan, M., Cao, M., Wang, Q., Li, Y., Yuan, G., Lei, M., and Huang, J. (2019). Structural basis of the crosstalk between histone H2B monoubiquitination and H3 lysine 79 methylation on nucleosome. Cell Res. 29, 330–333.

Zhang, J., Liang, Y., and Zhang, Y. (2011). Atomic-Level Protein Structure Refinement Using Fragment-Guided Molecular Dynamics Conformation Sampling. Structure 19, 1784–1795.

Zhang, Y. (2008). I-TASSER server for protein 3D structure prediction. BMC bioinformatics 9, 40.

Zheng, S.Q., Palovcak, E., Armache, J.P., Verba, K.A., Cheng, Y., and Agard, D.A. (2017). MotionCor2: anisotropic correction of beam-induced motion for improved cryo-electron microscopy. Nat. Methods 14, 331–332.

Zheng, W., Zhang, C., Wuyun, Q., Pearce, R., Li, Y., and Zhang, Y. (2019). LOMETS2: improved meta-threading server for fold-recognition and structure-based function annotation for distant-homology proteins. Nucleic Acids Res. 47(W1):W429–W436.

Zhou, B.R., Jiang, J., Feng, H., Ghirlando, R., Xiao, T.S., and Bai, Y. (2015). Structural Mechanisms of Nucleosome Recognition by Linker Histones. Mol. Cell 59, 628–638.

Zhou, K., Gaullier, G., and Luger, K. (2019). Nucleosome structure and dynamics are coming of age. Nat. Struct. Mol. Biol. 26, 3–13.

Zivanov, J., Nakane, T., Forsberg, B.O., Kimanius, D., Hagen, W.J., Lindahl, E., and Scheres, S.H. (2018). New tools for automated high-resolution cryo-EM structure determination in RELION-3. Elife 7. doi: 10.7554/eLife.42166.

