## Supplemental information for "Cryo-EM structure of the human Mixed Lineage Leukemia-1 complex bound to the nucleosome"

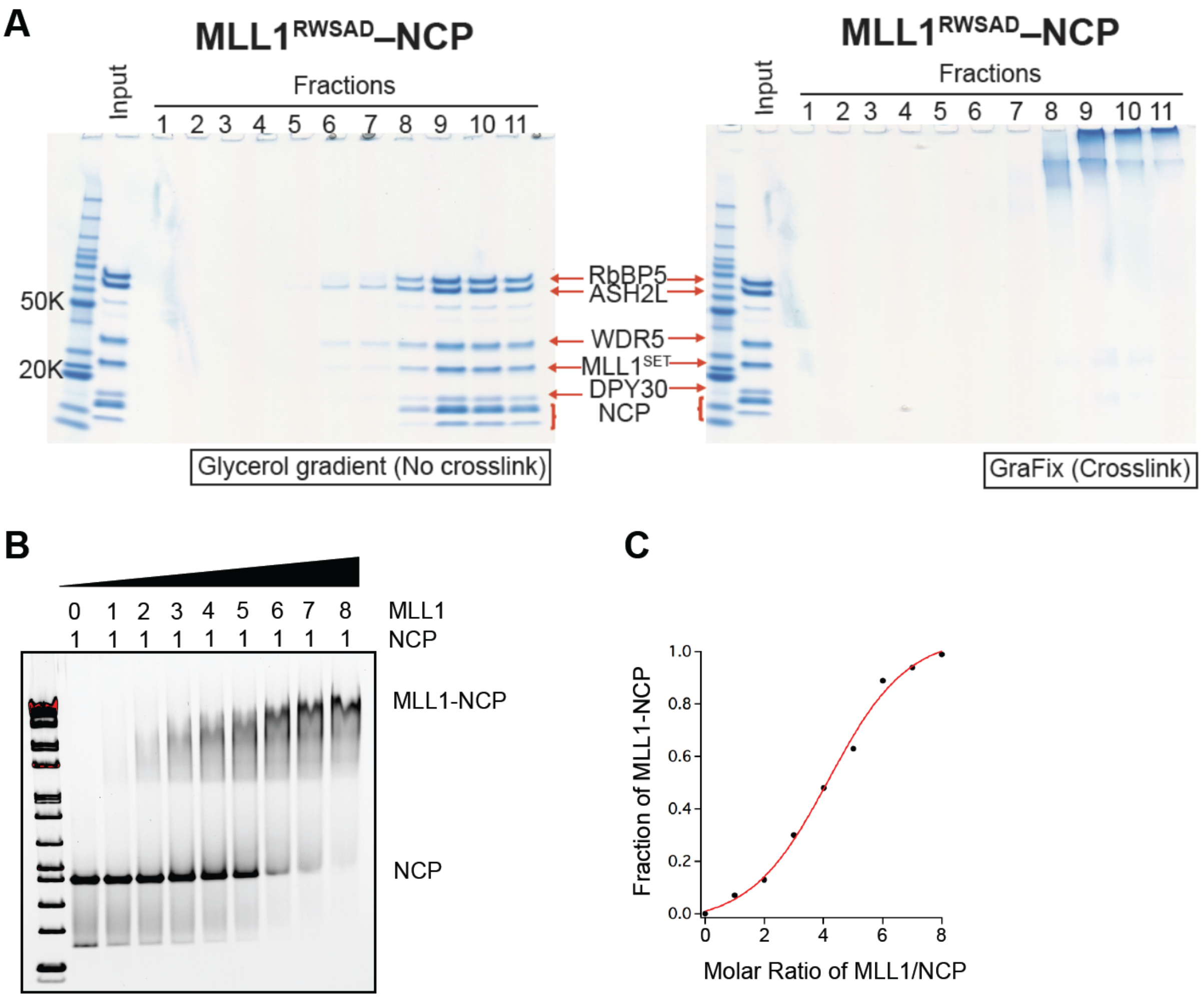


**Figure S1. Preparation of the MLL1$^{RWSAD}$-NCP Complex. This figure is related to Figure 1.**

(A) MLL1$^{RWSAD}$ complex and the NCP were incubated and isolated by glycerol-gradient (0-60%) with (right) or without (left) crosslinking. The fractions (1-11) were analyzed by SDS-PAGE. Individual components of the MLL1 core complex and the NCP were indicated.

(B) Gel mobility shift assay for the MLL1 core complex and the NCP. The molar ration of MLL1 vs. NCP was indicated on top.

(C) Quantification of the MLL1-NCP interaction with increasing concentration of the MLL1 core complex. It showed that MLL1-NCP interaction had modest affinity.

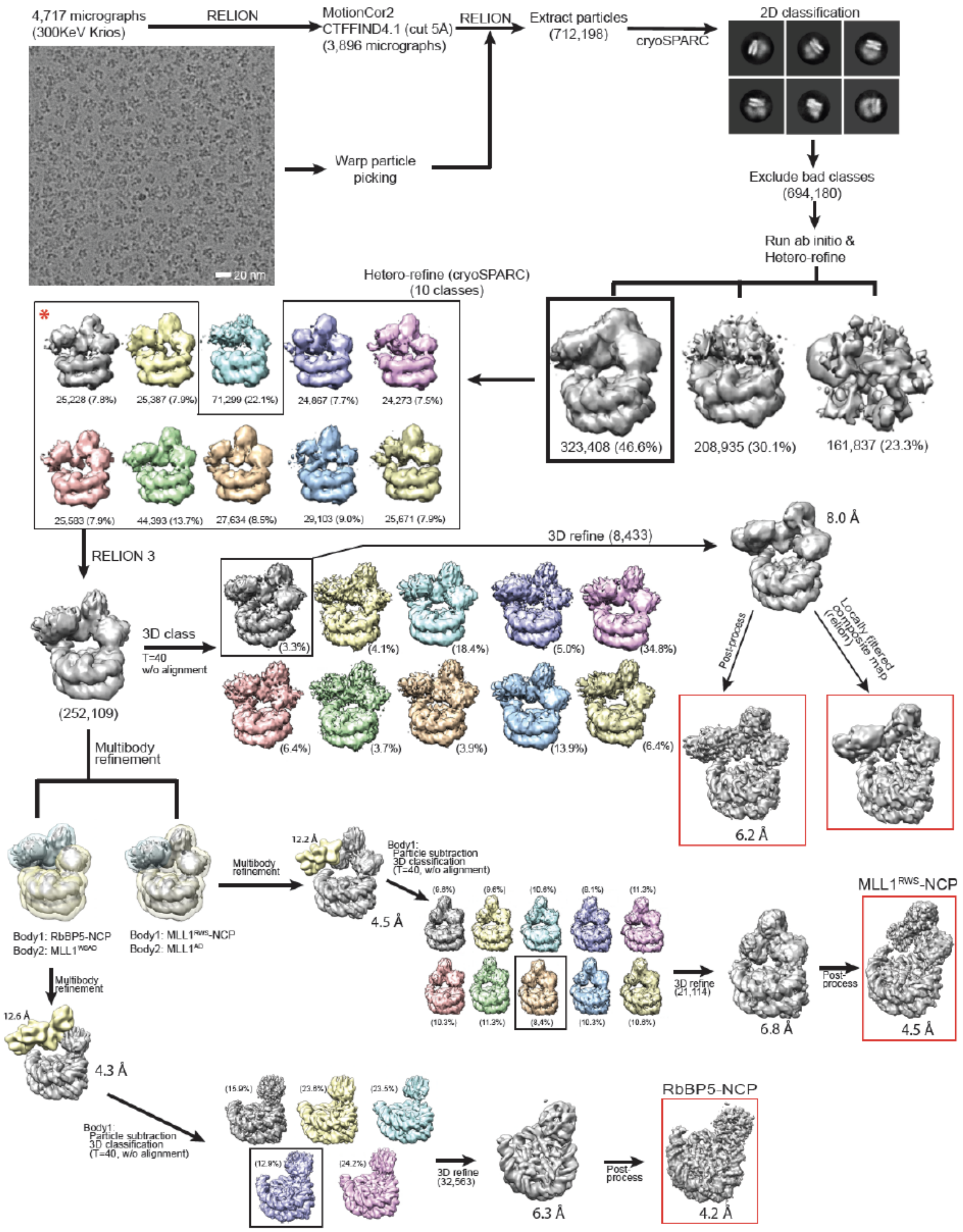


**Figure S2. Cryo-EM Data Processing for the MLL1$^{RWSAD}$-NCP. This figure is related to Figure 1 and Table 1.**

Representative micrograph image (Titan Krios 300 KeV) and 2D classifications of the MLL1$^{RWSAD}$-NCP complex were shown. The particle numbers for each classification step as well as the estimated resolution of overall and selected subcomplexes (red box) were shown at the bottom. The detailed data processing procedures of MLL1$^{RWSAD}$-NCP, MLL1$^{RWS}$-NCP, and RbBP5-NCP were described in the STAR methods.

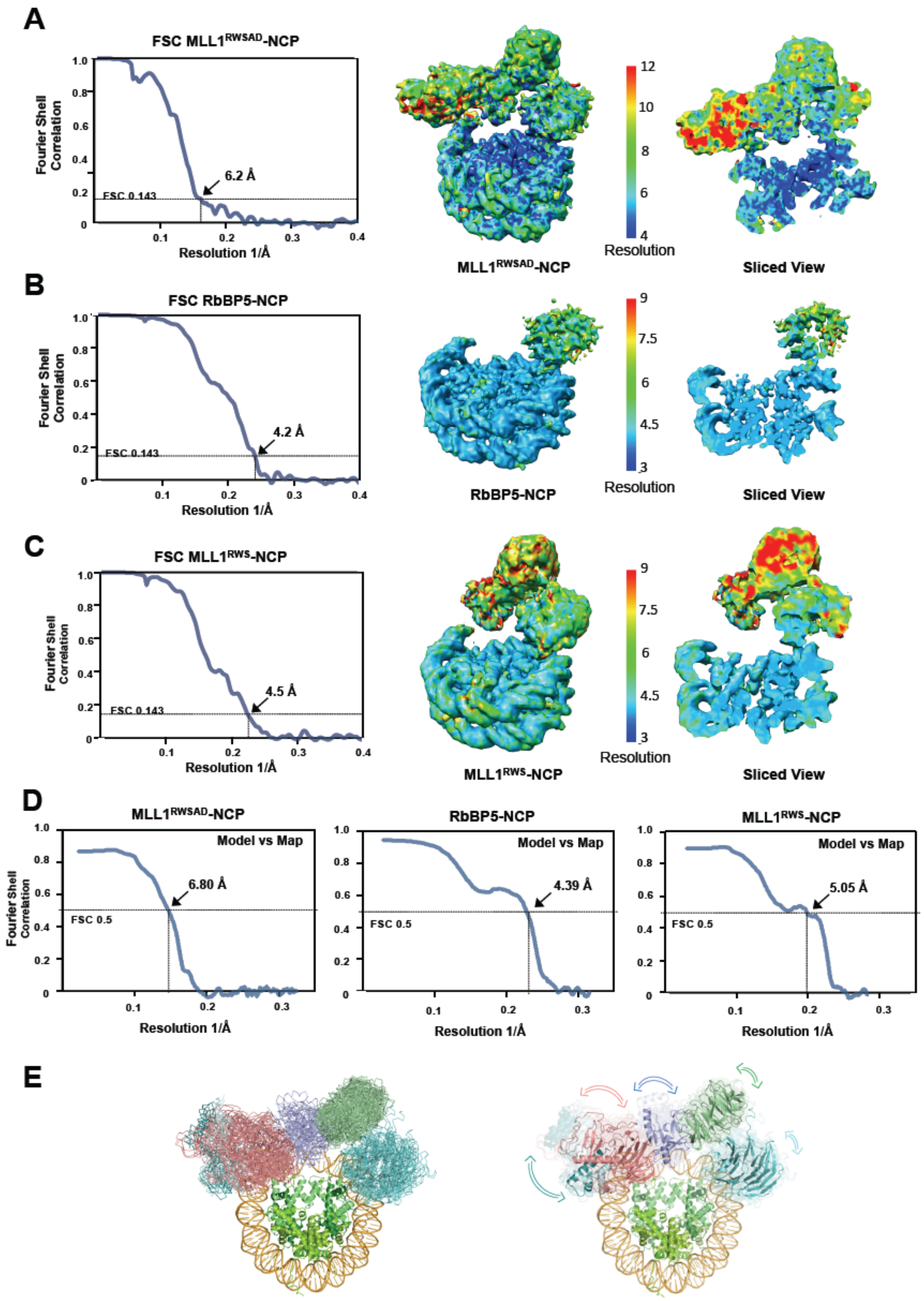


**Figure S3. Cryo-EM Map Validation of MLL1$^{RWSAD}$-NCP, RbBP5-NCP, and MLL1$^{RWS}$-NCP. This figure is related to Figure 1 and Table 1.**

(A-C) Fourier Shell Correlation (FSC) curves for MLL1$^{RWSAD}$-NCP (A), RbBP5-NCP (B), and MLL1$^{RWS}$-NCP (C) were shown on left and the corresponding local resolution assessments by RESMAP (Kucukelbir et al., 2014) were shown on right. The final resolution was determined using FSC=0.143 criterion, which was shown by arrowhead on the FSC curve.

(D) Model-map FSC curves for MLL1$^{RWSAD}$-NCP, RbBP5-NCP, and MLL1$^{RWS}$-NCP calculated by phenix.mtriage (Afonine et al., 2018). The resolution was indicated using FSC=0.5 criterion, which was shown by arrowhead on the FSC curve.

(E) Rigid body fitting of the MLL1$^{RWSAD}$ domains into 9 cryo-EM maps of MLL1$^{RWSAD}$-NCP, which were shown as hetero-refine subclasses in Figure S2 (highlighted by *). Left, nine coordinates of the MLL1$^{RWSAD}$ core complex were overlaid and displayed. Right, the degree of relative movement of each MLL1$^{RWSAD}$ domain within nine coordinates was indicated by arrows. The length and orientation of each arrow indicated the degree of dynamics and moving direction of each domain.

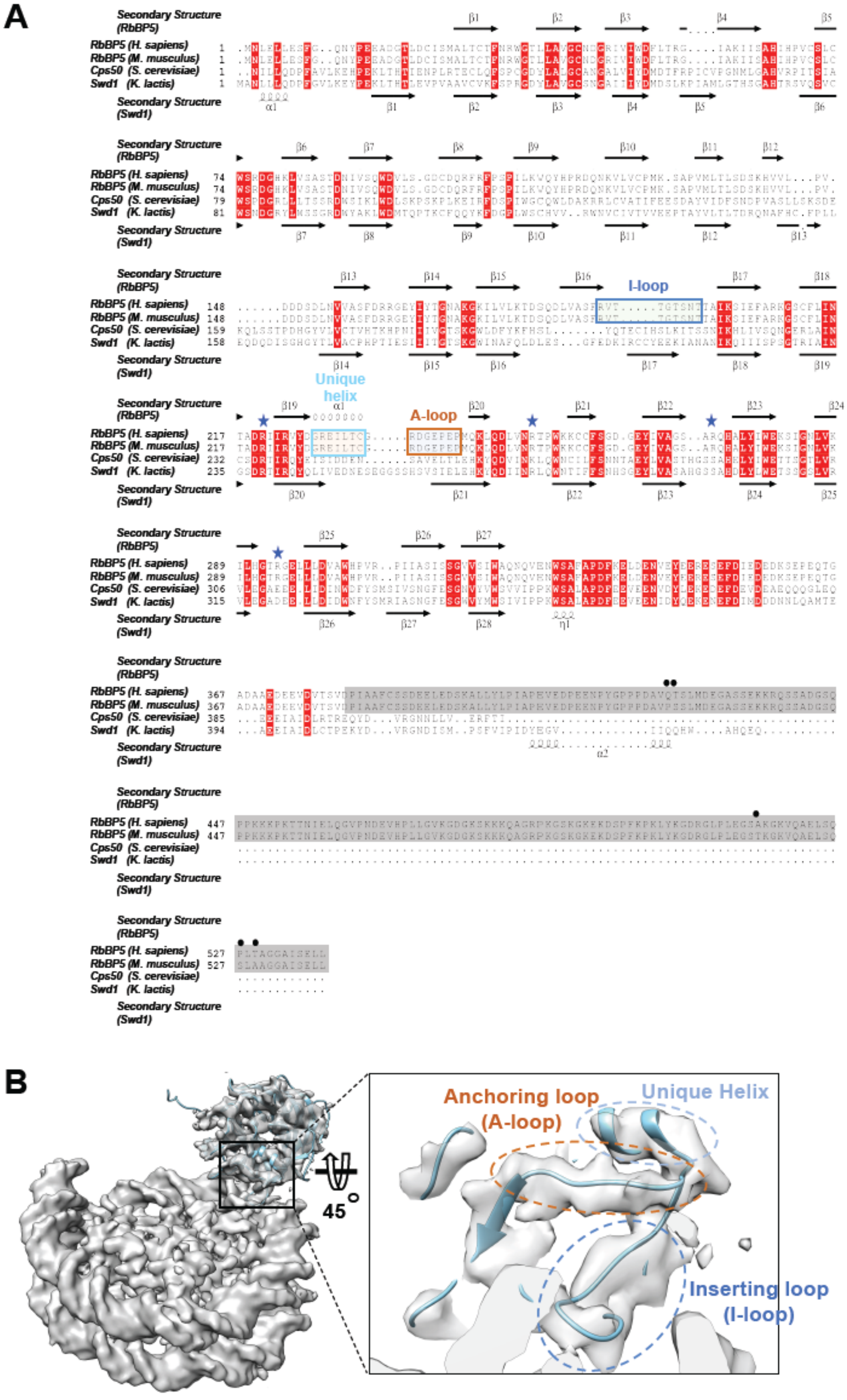


**Figure S4. The cryo-EM Structure of the RbBP5-NCP Subcomplex. This figure is related to Figure2.**

(A) The primary sequences of RbBP5 in *H. sapiens* and *M. musculus* (mammalian cells) as well as yeast homologous Swd1/Cps50 in *S. cerevisiae* and *K. lactis* were used for multiple sequence alignment. The secondary structures of mouse RbBP5 and Swd1 based on determined crystal structures were indicated on the top and bottom of the alignment, respectively. The I- and A-loops as well as the unique helix in mammalian RbBP5 were highlighted in blue, cyan, and orange boxes, respectively. Quad-R residues were shown as blue stars. Human and mouse RbBP5 had sequence divergence at five residues at C-terminus, which were indicated by black dots. The structural part of mouse RbBP5 WD40 repeats, which covers residues 1–380, are identical between human and mouse (100% sequence identity). RbBP5 C-terminus is not included in the crystal structure (grey box).

(B) Rigid-body fitting of RbBP5 (PDB ID: 5OV3) in the RbBP5-NCP cryo-EM map. Characteristic I- and A-loops as well as the unique helix (zoomed in left) of RbBP5 were fitted nicely into the cryo-EM map.

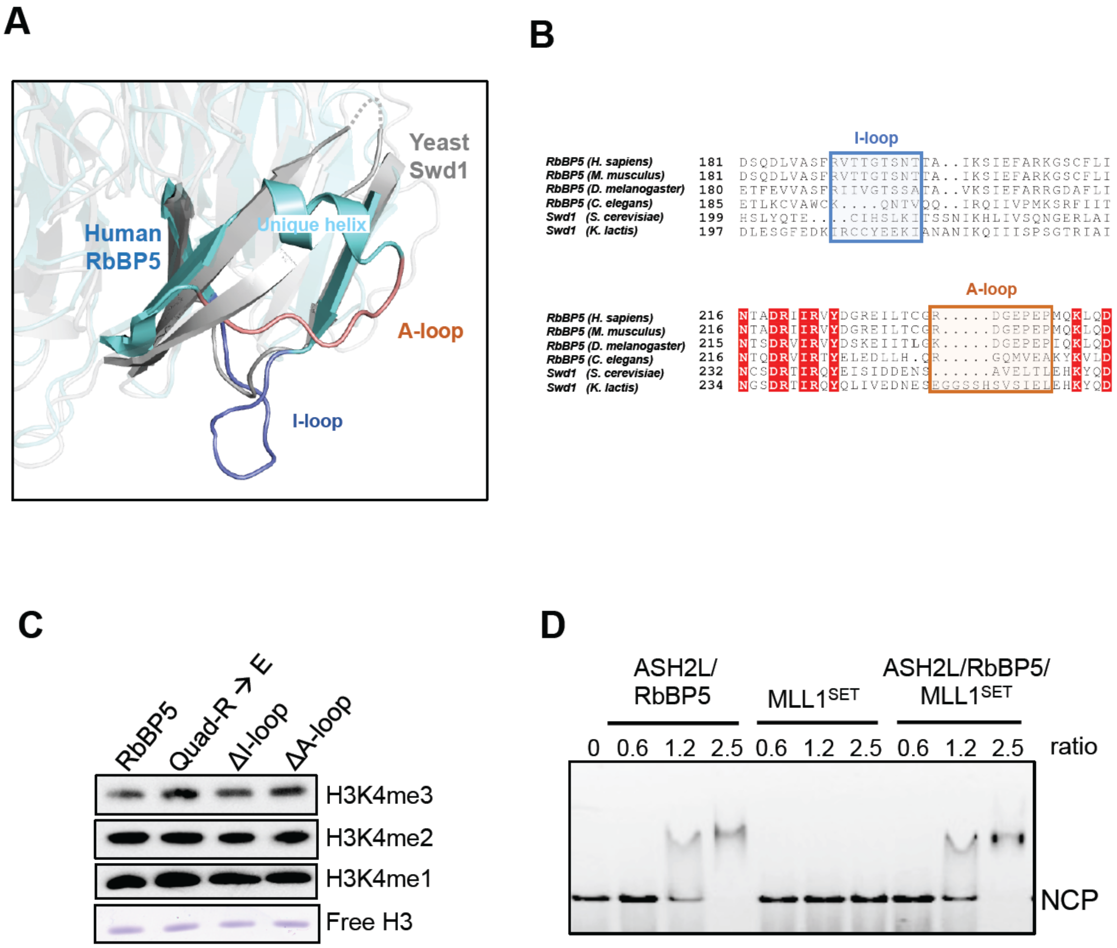


**Figure S5. Conservation and Divergence of RbBP5-NCP Interaction and Functional Implication. This figure is related to Figures 2 and 3.**

(A) Structural overlay of human RbBP5 and yeast Swd1 (PDB ID: 6CHG). RbBP5 (cyan) and Swd1 (gray) had distinct features for I- and A-loops.

(B) Multiple sequence alignment of RbBP5 I- and A-loops in eukaryotes. I- (blue box) and A- (red box) loops were highly conserved in from *D. melanogaster* to mammals.

(C) Immunoblot to detect *in vitro* histone methyltransferase activity using free histone H3 as the substrate. Reconstituted $MLL1^{RWSAD}$ complexes containing wild type and mutant RbBP5 were used, as indicated on top. Coomassie stain of H3 was included as the control.

(D) Electrophoretic mobility shift assay for the NCP with or without binding of protein or protein complexes as indicated on top. Molar ratio of protein(s) to the NCP was indicated on top.

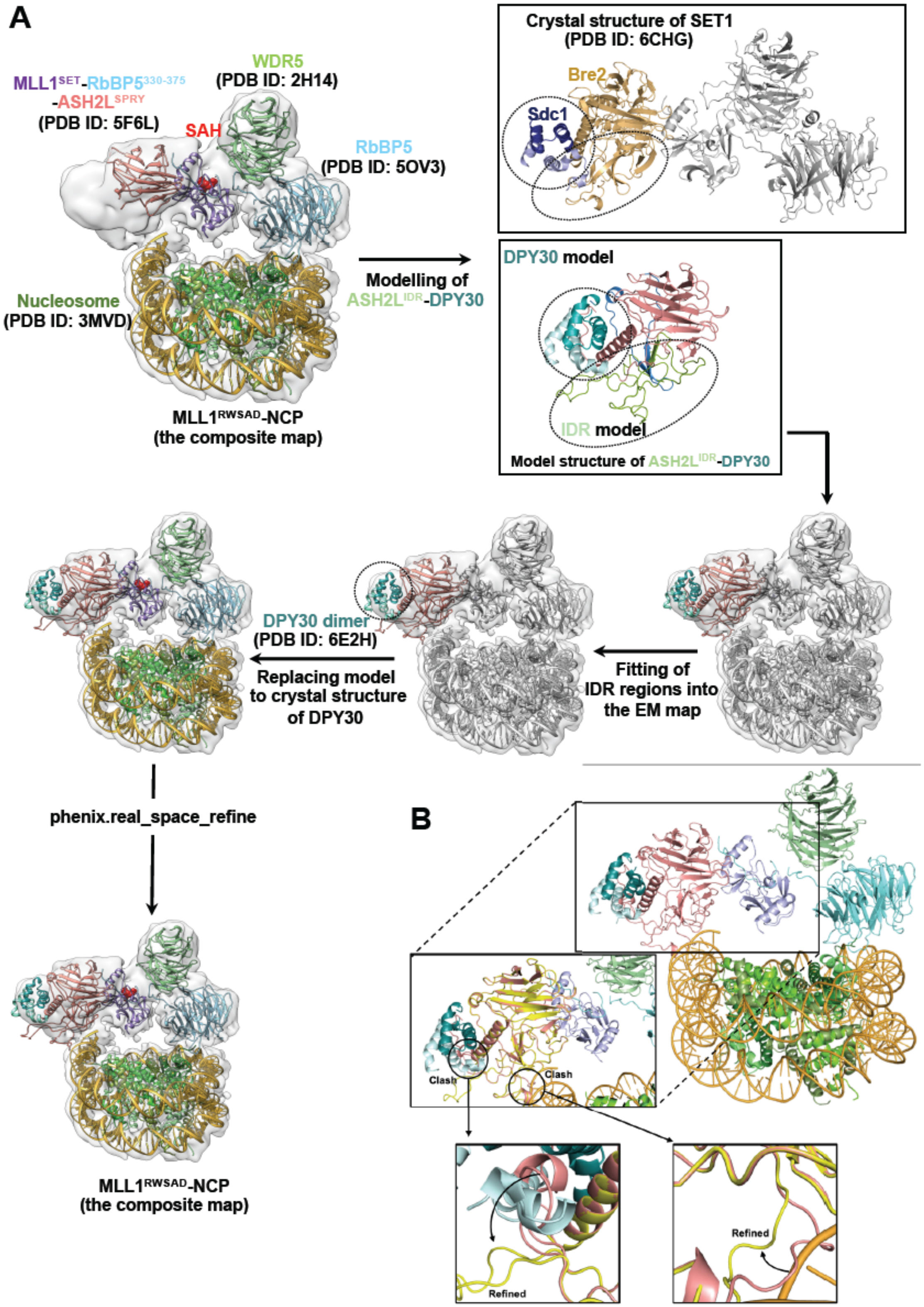


**Figure S6. Molecular Modeling of the ASH2L-NCP Interaction. This figure is related to Figure 4.**

(A) Flow chart of the molecular modeling of ASH2L-IDR (also see the STAR methods). The crystal structure of Bre2-Sdc1 from the yeast SET1 complex (PDB ID: 6CHG) was used as template. After modeling, DPY30 dimer was replaced by crystal structure of DPY30 dimer (PDB ID: 6E2H).

(B) Fragment-guided molecular dynamic refinement to remove two clashes (black circle; pink-before, yellow-after) between ASH2L-IDR and DPY30 or DNA using the software FG-MD (Zhang et al., 2011).

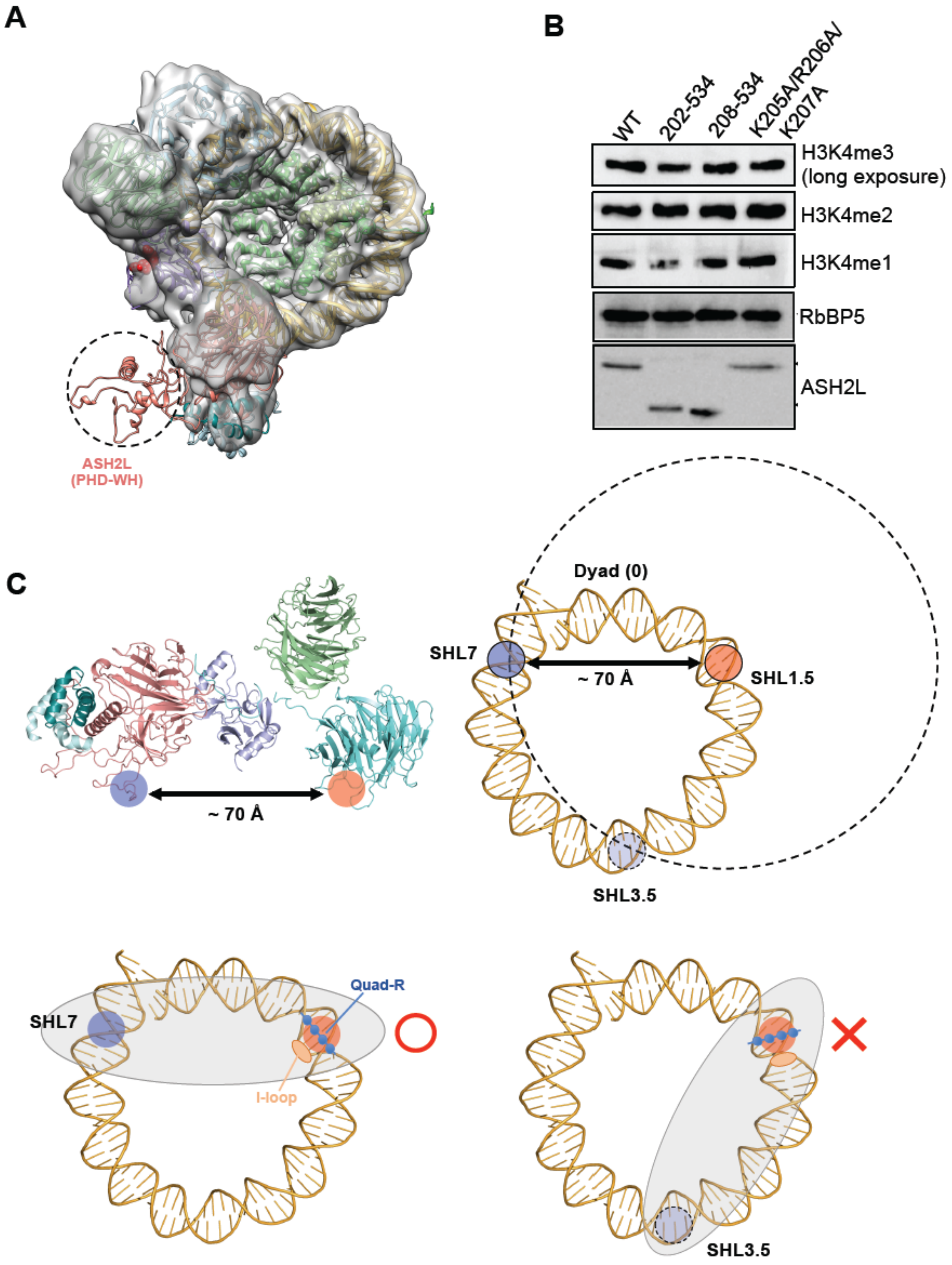


**Figure S7. ASH2L-NCP Interaction in the MLL1$^{RWSAD}$-NCP complex. This figure is related to Figure 5, 6 and 7.**

(A) The ASH2L PHD-WH domain (model structure; black dashed circle) was not visible in the cryo-EM map of MLL1$^{RWSAD}$-NCP. One potential position of the PHD-WH domain based on the structure prediction was shown.

(B) Immunoblot to detect *in vitro* histone methyltransferase activity using free histone H3 as the substrate. Reconstituted MLL1$^{RWSAD}$ complexes containing wild type and mutant ASH2L were used, as indicated on the top. Immunoblot for RbBP5 and ASH2L were included as controls.

(C) The model for coordinated binding to the NCP by RbBP5 and ASH2L. The distance between the I-loop of RbBP5 (red circle) and the '$_{205}$-KRK-$_{207}$' basic patch of ASH2L Linker-IDR (blue circle) is ~ 70 Å. Specific anchoring of RbBP5 on the NCP confers both orientation and distance constraints for ASH2L-NCP binding. The binding of ASH2L at DNA SHL3.5 may not be allowed due to unfavorable interactions of Quad-R/DNA and I-loop/H4 tail in RbBP5-NCP (bottom right).
